# Facilitatory stimulation of the pre-SMA enhances semantic cognition via remote network effects on task-based activity and connectivity

**DOI:** 10.1101/2022.10.21.513185

**Authors:** Sandra Martin, Regine Frieling, Dorothee Saur, Gesa Hartwigsen

**Author notes:** Correspondence: Sandra Martin.

## Abstract

**Background:** The continuous decline of executive abilities with age is mirrored by increased neural activity of domain-general networks during task processing. So far, it remains unclear how much domain-general networks contribute to domain-specific processes such as language when cognitive demands increase. The current neuroimaging study explored the potential of intermittent theta-burst stimulation (iTBS) over a domain-general hub to enhance executive and semantic processing in healthy middle-aged to older adults.

**Methods:** We implemented a cross-over within-subject study design with three task-based neuroimaging sessions per participant. Using an individualized stimulation approach, we stimulated each participant once with effective and once with sham iTBS over the pre-supplementary motor area (pre-SMA), a region of domain-general control. Subsequently, task-specific stimulation effects were assessed in functional MRI using a semantic and a non-verbal executive task with varying cognitive demand.

**Results:** Effective stimulation increased activation relative to sham stimulation only during semantic processing in visual and dorsal attention networks. Further, iTBS induced increased functional connectivity in task-specific networks for semantic and executive conditions with high cognitive load. Notably, stimulation-induced changes in activation and connectivity related differently to behavior: While increased activation of the parietal dorsal attention network was linked to poorer semantic performance, its enhanced coupling with the pre-SMA was associated with more efficient semantic processing.

**Conclusions:** iTBS modulates networks in a task-dependent manner and generates effects at regions remote to the stimulation site. These neural changes are linked to more efficient semantic processing, which underlines the general potential of network stimulation approaches in cognitive aging.

## Introduction

Non-invasive brain stimulation (NIBS) techniques are recognized as a promising approach to counteract age-related cognitive decline and to promote successful aging. These techniques may have the potential to support preservation of cognitive functions in pathological but also healthy aging through modulation of cortical excitability and the enhancement of neuroplasticity [1,2]. Especially the application of theta-burst stimulation (TBS), a patterned protocol of repetitive transcranial magnetic stimulation (rTMS), has gained increasing interest since it is thought to induce longer-lasting after-effects [3]. So far, only a few studies with heterogeneous results explored the modulatory effects of TBS on cognition in aging brains [4–7].

Further insight can be gained by investigating the effect of stimulation on neural activity and functional connectivity. Neuroimaging results can help interpreting behavioral effects and might even be observed in the absence of a stimulation-induced behavioral change [8]. A better understanding of the modulatory effects of TBS at the neural level would increase the efficiency of network stimulation in aging brains. Such network approaches may be more powerful than conventional modulatory applications that target specific brain regions within specialized networks [9].

The continuous decline of executive abilities is a hallmark of cognitive aging. Its effects can also be observed in domains that usually remain well preserved, such as language and creativity, when contextual demands are high [10,11]. For instance, in the domain of semantic cognition, the decline of cognitive control is reflected by poorer semantic selection processes with age, such as inhibiting irrelevant semantic associations, but not semantic representations per se [12].

On the neural level, age-related changes in semantic cognition are mirrored by a shifted network architecture with increased activity of domain-general networks during task processing [13] and increased coupling of distinct functional networks [14]. So far, it remains unclear how much domain-general networks, such as the multiple-demand network (MDN), contribute to the maintenance of semantic processing in healthy aging. Specifically, the neural network of semantic control may be multidimensional, consisting of domain-specific semantic control, which subserves processes such as the controlled retrieval of less salient conceptual features, and domain-general control, which supports general selection and inhibition mechanisms. This notion is supported by the observation that brain regions that are active in tasks with high semantic control demands partially overlap with areas of the MDN [15].

This study aimed to determine if facilitatory TBS could enhance semantic and executive task processing in healthy middle-aged to older adults. The pre-supplementary motor area (pre-SMA) was targeted for stimulation using effective and sham intermittent TBS (iTBS), as it is associated with the semantic control network and the domain-general MDN [15,16], and contributes more to semantic processing in older adults [14,17]. We included middle-aged to older adults since executive functions like alertness and inhibitory control have been shown to decline from middle age [18,19], which might increase the modulatory potential of TBS in this age group. Using a verbal semantic judgment task with varying cognitive demand and a non-verbal tone judgment task, we were interested in the effect of iTBS on task-based activity and functional connectivity and its relationship with behavior. Specifically, we implemented a cognitively more demanding feature-picture matching (FPM) and a low-level word-picture matching (WPM) task to distinguish stimulation effects on semantic control. Further, the tone judgment task was included to characterize the modulatory effect of pre-SMA stimulation on non-verbal executive demands.

We hypothesized that iTBS modulates processing efficiency through faster reactions on the behavioral level and increased whole-brain activity as well as enhanced remote network modulation for our conditions with high cognitive load, FPM and tone judgment. Further, if the pre-SMA contributes to semantic control, comparing FPM with the low-level WPM might reveal task-specific stimulation effects beyond general cognitive control. Finally, we aimed to elucidate how stimulation-induced changes in activity and connectivity relate to behavioral modulation.

## Material and methods

### Participants

Thirty healthy middle-aged to older adults (14 female; *M* = 61.6, *SD* = 7.64, range: 45–74 years) participated in the experiment. Inclusion criteria were native German speaker, right-handedness, normal hearing, normal or corrected-to-normal vision, no history of neurological or psychiatric conditions, and no contraindication to MRI or rTMS. Participants were also screened for cognitive impairments using the Mini-Mental-State-Examination [20] (all ≥ 26/30 points). Written informed consent was obtained from each participant. The study was approved by the local ethics committee and conducted in accordance with the Declaration of Helsinki.

### Experimental Design

Figure 1 displays the experimental procedure. We employed a single-blind, cross-over design with three sessions per participant (Figure 1A). Sessions were separated by at least one week (mean inter-session interval: 28.4 days; *SD*: 51.2). During the first session (baseline), participants completed two runs of a language localizer task and two runs of the experimental task. The second and third session each began with effective or sham iTBS over the pre-SMA (Figure 1C). Participants subsequently completed two runs of the experimental paradigm. The order of effective and sham sessions was counterbalanced across participants.

**Figure 1.**
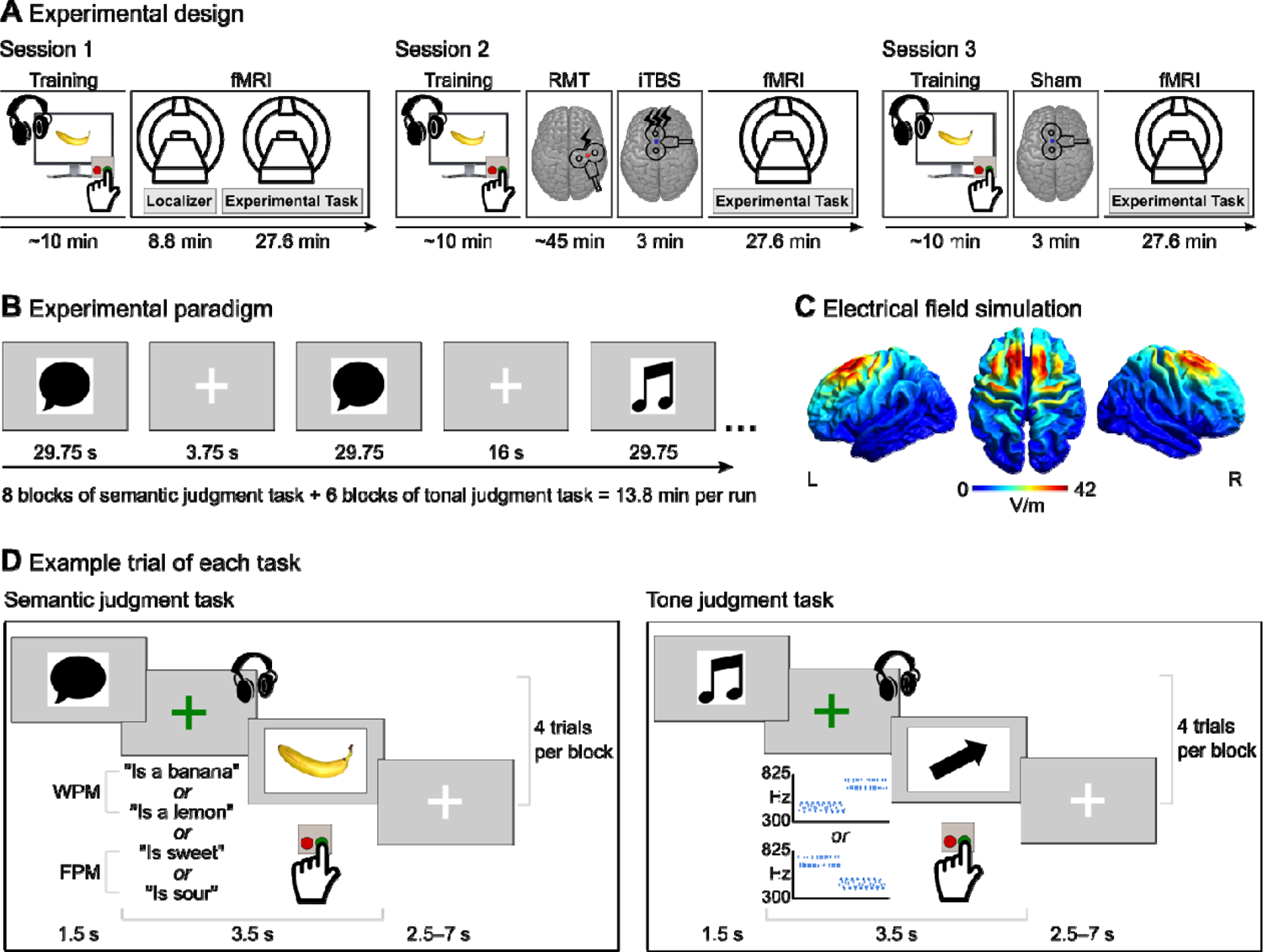
Experimental Design. (A) Participants completed three sessions: a baseline fMRI session and two iTBS + fMRI sessions with effective and sham stimulation. (B) Per fMRI session, two task runs were completed. Blocks of the semantic judgment and the tone judgment task were interspersed with rest blocks. (C) We simulated the average electrical field at our target site, the pre-SMA (visualized on the normalized cortical surface). (D) Example trials for the semantic and the tone judgment task are shown. Participants heard a short phrase or a sequence of two tones. At the offset of the auditory stimulus, a picture of an object or an arrow appeared. Participants indicated via button press whether auditory and visual stimuli matched. RMT: resting motor threshold, WPM: word-picture matching, FPM: feature-picture matching.

### Experimental Paradigm

Two tasks were implemented in the fMRI experiment: a semantic judgment task with varying cognitive demand (low demand: WPM, high demand: FPM) and a non-verbal tone judgment task (Figure 1B and D). Stimuli of the tasks are described in more detail in Supplementary Methods. In both tasks, participants were required to decide whether an auditory stimulus matches with a presented image via yes/no-button press using the index and middle finger of their left hand. The left hand was used to shift motor activity related to the button press to the right hemisphere. The order of buttons was counterbalanced across participants. Tasks were presented in mini-blocks of four trials per task and blocks were separated by short rest intervals. Individual trials were 3.5 s long including presentation of auditory and visual stimulus, and button press by the participant. Each run included 88 stimuli with 32 items per condition of semantic judgment and 24 items of tone judgment. Participants completed two runs per session.

### Magnetic Resonance Imaging

MRI data were collected at a 3T Siemens Magnetom Skyra scanner (Siemens, Erlangen, Germany) with a 32-channel head coil. For functional scans, a gradient-echo echo-planar imaging multiband sequence [21] was used with the following parameters: TR: 2000 ms, TE: 22 ms, flip angle: 80°, voxel size: 2.48 x 2.48 x 2.75 mm, FOV: 204 mm, multiband acceleration factor: 3, 60 axial whole-brain slices with interleaved order. For the language localizer task, 266 volumes were acquired. For the experimental task, a total of 842 volumes per session were acquired. For distortion correction, field maps (pepolar images) were obtained at the end of each session (TR: 8000 ms, TE: 50 ms). A T1-weighted volume was acquired using an MPRAGE sequence (176 slices, whole-brain coverage, TR: 2300 ms, TE: 2.98 ms, voxel size: 1 x 1 x 1 mm, matrix size: 256 x 240 mm, flip angle: 9°).

### Intermittent Theta Burst Stimulation

rTMS was delivered using the iTBS protocol which consists of bursts of three pulses at 50 Hz given every 200 ms in two second trains, repeated every ten seconds over 190 s for a total of 600 pulses [22]. We chose TBS since its high-frequency protocols have been reported to induce longer lasting after-effects with a duration of up to one hour [3]. iTBS was applied via stereotactic neuronavigation (TMS Navigator, Localite, Bonn, Germany) and a MagPro X100 stimulator (MagVenture, Farum, Denmark) with an MCF-B65 figure-of-eight coil. For sham stimulation, we used the corresponding placebo coil. During stimulation, the coil handle was pointed in a 90° angle [e.g., 23,24] and intensity was set to 90% of individual resting motor threshold (RMT). RMT was defined as the lowest stimulation intensity producing at least five motor evoked potentials of ≥50 µV in the relaxed first dorsal interosseous muscle when single-pulse TMS was applied to the right motor cortex ten times.

Stimulation coordinates for iTBS over the pre-SMA were based on individual activation patterns for the contrast semantic judgment > rest within a pre-defined region of interest (ROI) mask of the pre-SMA [25]. The stimulation target was defined as the global peak of the strongest cluster within the pre-SMA ROI in individuals’ subject space after FWE-correction at *p* < 0.05. For plots of the individual stimulation sites within the ROI, see Figures S1 and S2. We performed electrical field simulations using SimNIBS v.4.0.0 [26] to characterize location, extent, and strength of the electrical field induced by iTBS over the pre-SMA in each individual subject. Figure 1C displays the average electrical field while individual fields are shown in Figure S3.

### Data Analyses

#### Behavioral data

Accuracy and reaction time data of each session were analyzed using mixed-effects models with a logistic regression for accuracy data due to their binary nature and a linear regression for log-transformed reaction time data. We only analyzed reaction times for correct responses. Contrast coding was done via sum coding. Based on our research questions, session (i.e., baseline, effective or sham stimulation) and condition (WPM, FPM or tone judgment) along with their interaction term were always entered as fixed effects. Next, we used stepwise model selection to determine the best-fitting model based on the Akaike Information Criterion. Tables S1 and S2 display the model selection procedures. Equations 1 and 2 show the best-fitting model for accuracy and reaction time data, respectively. Statistical models were performed with R v.4.2.1 [27] and the packages lme4 [28] for mixed models and bblme [29] for model comparisons. Plots and result tables were generated using the packages sjPlot [30] and ggeffects [31].

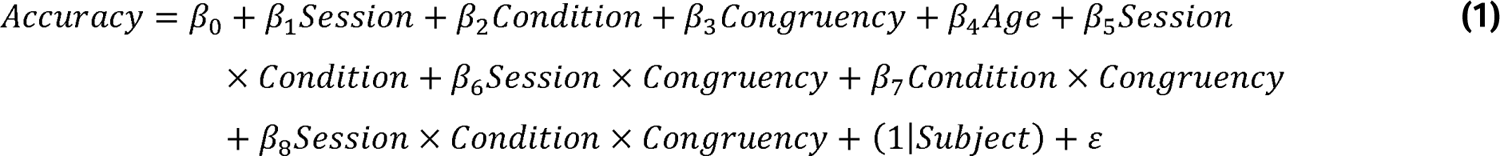

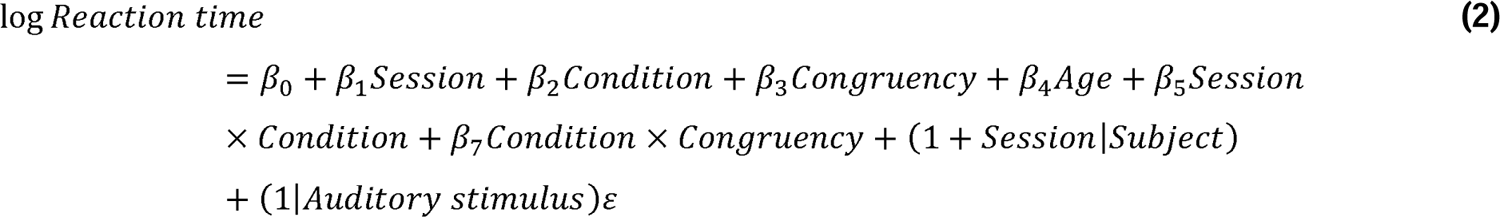

#### Whole-brain analyses

Preprocessing of MRI data was performed using fMRIPrep 20.2.3 [32] which is based on Nipype 1.6.1 [33]. A detailed description can be found in Supplementary Materials. Functional MRI data were modelled using a general linear model (GLM) for each session and participant. For the localizer, the GLM included regressors for the task blocks of intact and degraded listening. For the experimental task, regressors for the three conditions and a separate regressor for error trials were included in the GLM. To account for condition- and trial-specific differences in reaction time, the duration of a trial was defined as the length of the auditory stimulus plus the reaction time. Additional regressors included six motion parameters and individual regressors for strong volume-to-volume movement as indicated by values of framewise displacement > 0.7. Further, temporal and spatial derivatives were modelled for each condition, and a high-pass filter was applied to remove low-frequency noise.

Contrast images were entered intro group-level random effects models. For the first session, one-sample t-tests were computed to define condition-specific activation. To assess differences between effective and sham iTBS, contrast images from the sham session were subtracted from the effective session, and the difference images were submitted to random effects models where session effects were estimated using one-sample t-tests. Results were thresholded at p < 0.05 at peak level and corrected at cluster level for FWE rate at p < 0.05. We also tested for a potential effect of age on fMRI results and included age as covariate in group results. Since we did not detect significant effects of age, subsequent results are reported without this confound.

To assess the relationship between differences in activation and differences in behavior induced by iTBS, we extracted percent signal change (PSC) for our a-priori defined stimulation site pre-SMA and for clusters showing a significant effect of stimulation (*n* = 5; see Table 1) using the MarsBar toolbox [34]. For the pre-SMA, PSC was extracted for a cluster centered at each individual stimulation site and containing the 25% strongest activated voxels for the contrast semantic judgment > rest, which was identical to the contrast used for the definition of the stimulation site. Data were then entered into correlation analyses where the difference in PSC for a certain condition was correlated with the difference in accuracy and reaction time between effective and sham sessions.

**Table 1.**
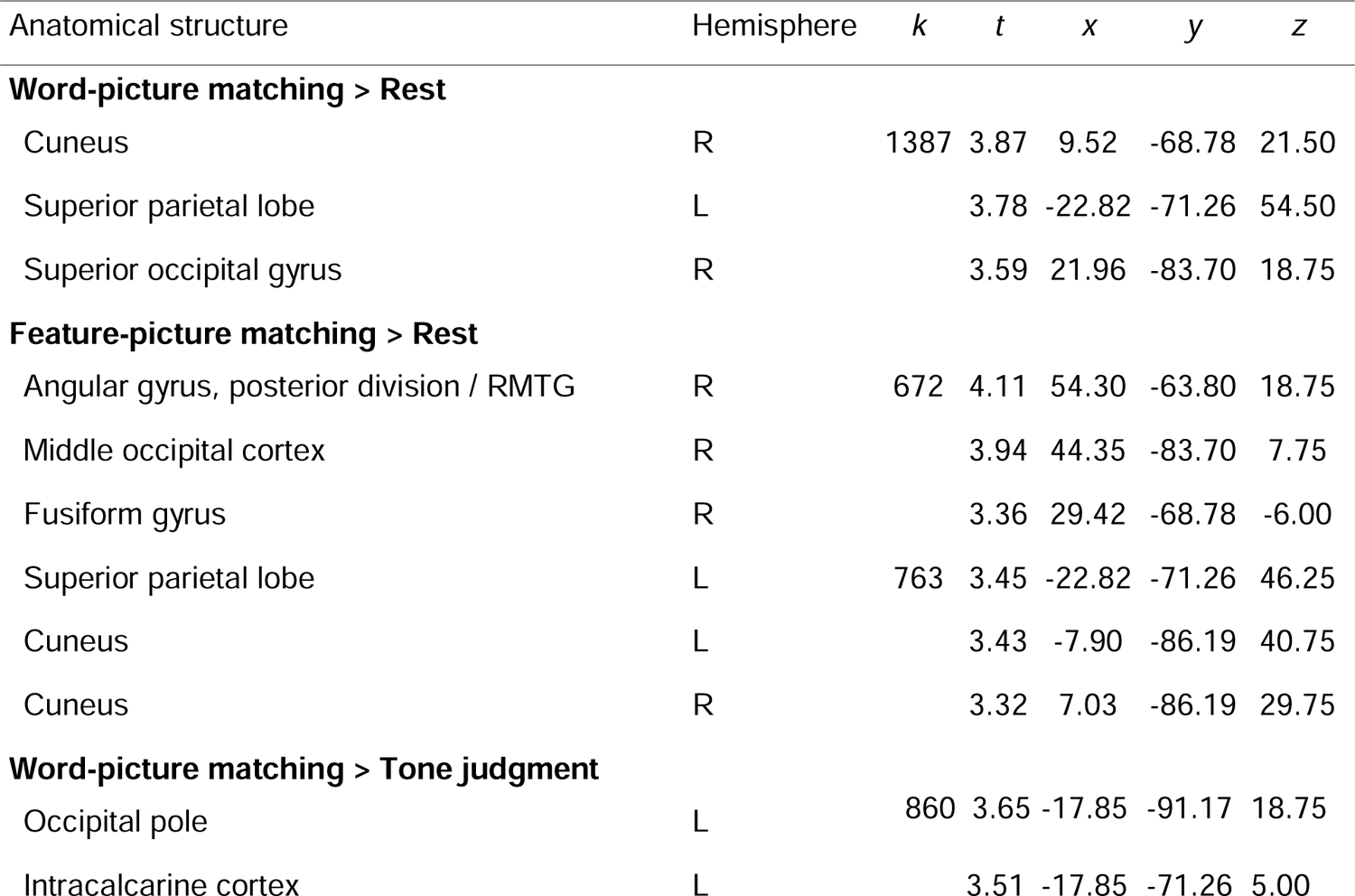
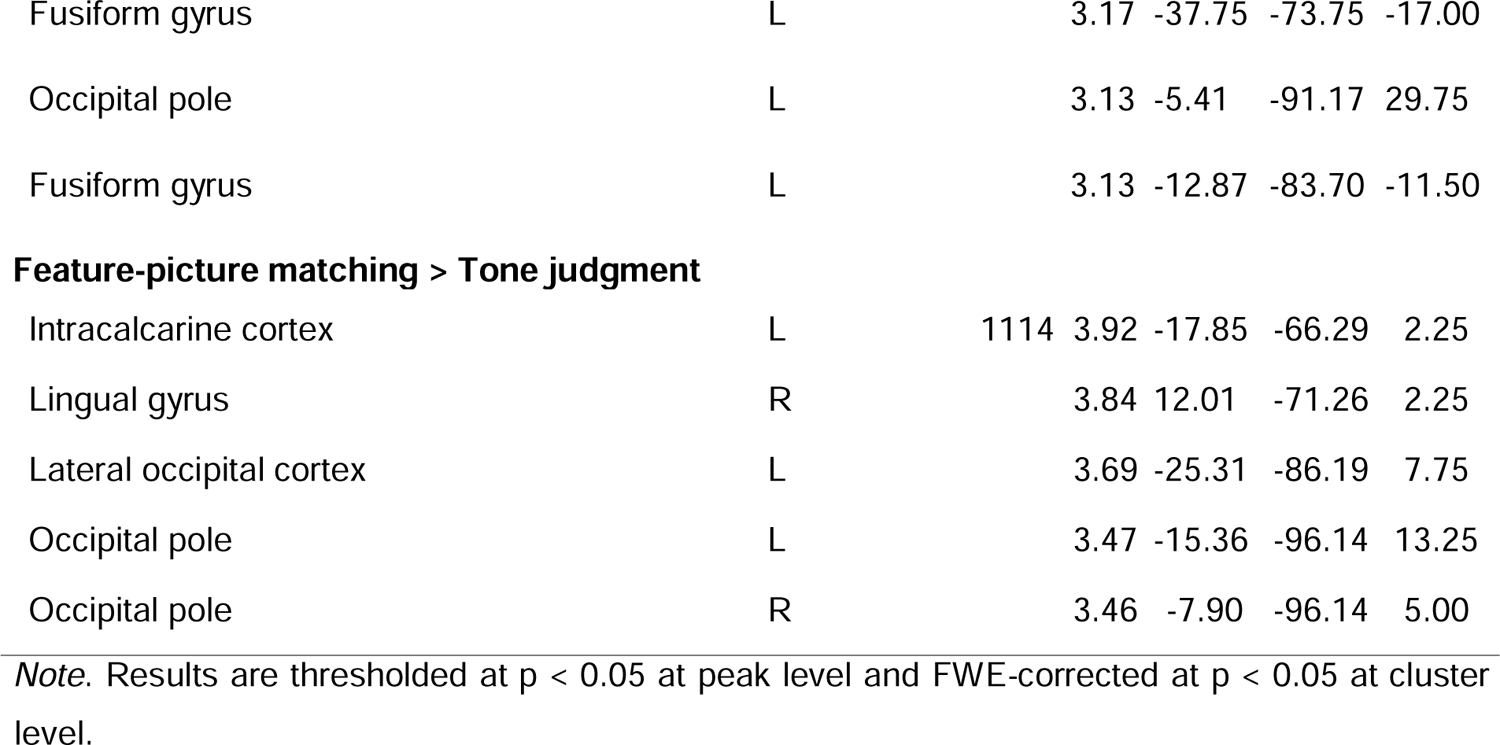
Univariate fMRI results – effective > sham iTBS

#### Analysis of subject-specific functional regions of interest

Data from the language localizer task were analyzed employing the group-constrained subject-specific approach [35]. This method allows the identification of individual functional ROIs sensitive to language processing [36], which were then used to characterize response profiles in the independent data set of the experimental task. We did not find any significant effect of stimulation session on activation in the functional ROIs. Details on analysis and results of subject-specific functional ROIs are described in Supplementary Methods and visualized in Figures S4-7.

#### Functional connectivity analysis

To assess potential changes in functional connectivity induced by iTBS, we conducted psychophysiological interaction (PPI) analyses using the gPPI toolbox [37]. Seed regions were defined for significant global cluster peaks for the contrast of effective and sham session (*n* = 5, cf. Table 1) and for our stimulation site, bilateral pre-SMA. Binary, resampled masks were created for each seed by building a spherical ROI with a radius of 10 mm. Next, individual ROIs were built by extracting the 25% most active voxels in each seed mask of a given contrast image.

For the gPPI, individual regression models were set up for each ROI and session containing the deconvolved time series of the first eigenvariate of the BOLD signal from the respective ROI as the physiological variable, regressors for the three task conditions and errors as the psychological variable, and the interaction of both variables as the PPI term. Models were adjusted for an omnibus F-test of all task regressors. Subsequently, first-level GLMs were calculated. We were specifically interested in potential differences between effective and sham iTBS sessions for the contrasts FPM > WPM and FPM > tone judgment. Random-effects models for group analysis were set up as described for the univariate analysis. Results were thresholded at p <0.01 at peak level and FWE-corrected p <0.05 at cluster level.

We also explored a relationship between stimulation-induced changes in functional connectivity and behavior. To this end, we extracted pre-SMA-to-ROI PPI connectivity for effective and sham sessions for the contrast semantic judgment > tone judgment where ROI refers to the five seed regions described above. We then correlated the difference between effective and sham connectivity for each pre-SMA-ROI pair with the difference between effective and sham in accuracy and reaction time.

## Results

### Behavioral results

For accuracy, the three-way interaction between session, condition, and congruency was not significant (*x*^2^= 7.83, *p* = 0.099). However, we detected a significant interaction between condition and congruency (*x*^2^= 53.15, *p* < 0.001) and session and condition (*x*^2^= 21.8, *p* < 0.001; Figure 3C). Post-hoc tests showed that incongruent items had higher accuracy in semantic conditions (WPM: *OR* = 0.38, *p* < 0.001; FPM: *OR* = 0.31, *p* < 0.001) but not tone judgments (*OR* = 1.18, *p* = 0.25). For session and condition, post-hoc tests showed a significant difference only for the tone task, such that participants performed generally better after the baseline session (active iTBS > baseline: *OR* = 0.41, *p* < 0.001; sham iTBS > baseline: *OR* = 0.57, *p* = 0.002). Moreover, we found main effects of condition (*x*^2^= 279.34, *p* < 0.001) and congruency (*x*^2^= 77.32, *p* < 0.001) but not of session (*x*^2^= 2.79, *p* = 0.25). Post-hoc tests revealed overall better performance for WPM than FPM (*OR* = 5.77, *p* < 0.001) and tone judgment (*OR* = 3.55, *p* < 0.001). For congruency, accuracy was higher for incongruent than congruent items (*OR* = 0.5, *p* < 0.001).

**Figure 3.**
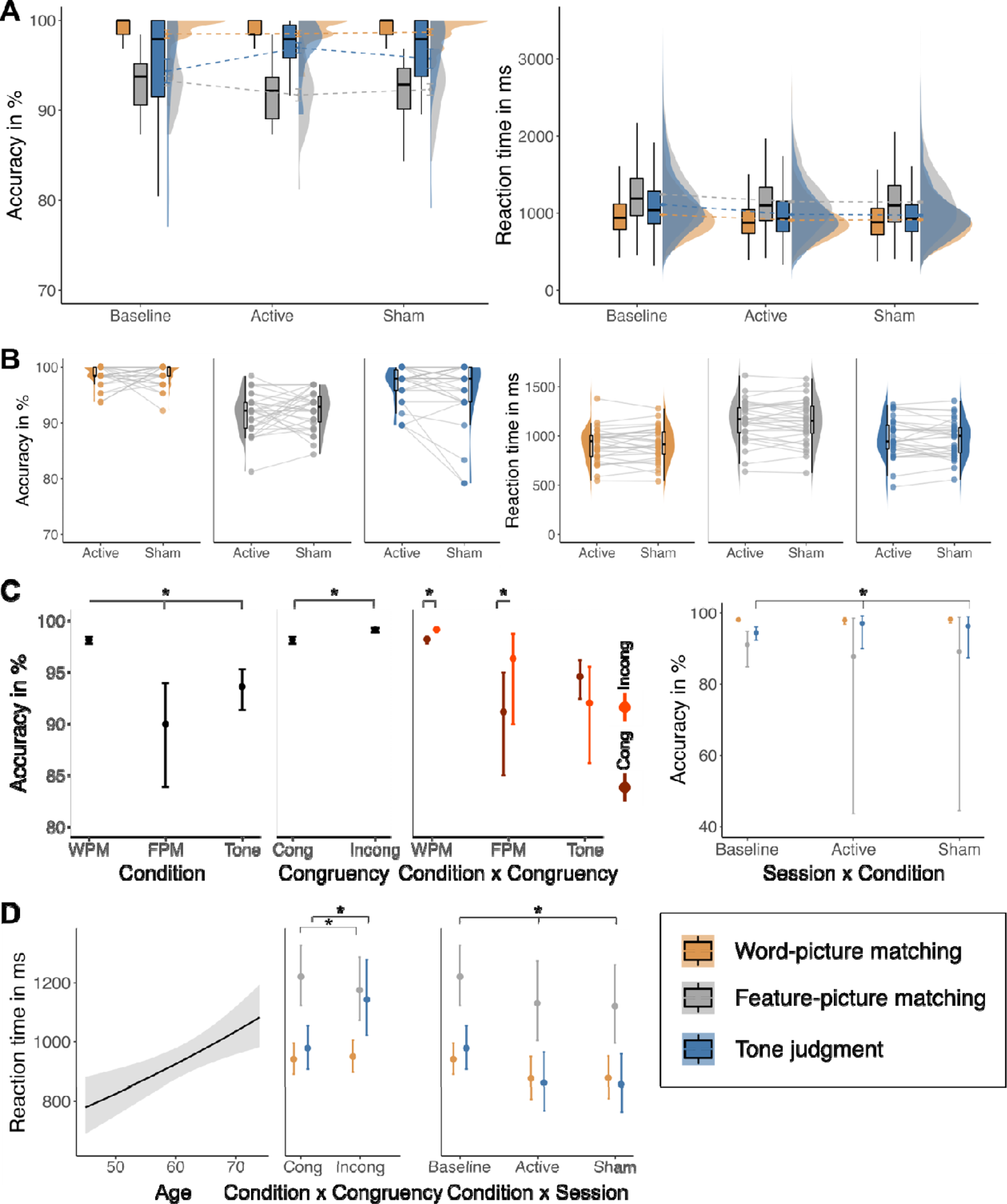
Behavioral results. (A) Results for accuracy and reaction time for each condition at each session. Boxplots show median and 1.5 x interquartile ranges. Half-violin plots display distribution and dotted lines show changes of mean values across sessions. (B) Individual data for effect of stimulation sessions on accuracy and reaction time for each condition. Results of mixed-effects regression (predicted marginal effects) for (C) accuracy and (D) reaction time. Full results output of both models can be found in Table S4. Cong: congruent items, Incong: incongruent congruent items were faster (*p* < 0.001). Further, we found that reactions times generally increased with age (*x*^2^= 9.4, *p* = 0.002). Full results of both models are shown in Supplementary Table S4.

For reaction time, results showed a significant interaction of session with condition (*x*^2^= 44.4, *p* < 0.001; Figure 3D). Reaction times improved for all three conditions after the baseline session (all *p* < 0.01). However, there was no difference in reaction time between effective and sham iTBS sessions. Results also showed a significant interaction between condition and congruency (*x*^2^= 306.53, *p* < 0.001). For FPM, incongruent items were faster (*p* = 0.034), while for tone judgment, items.

### Univariate functional MRI data

#### The effect of conditions at baseline

Both semantic and tone judgment activated large whole-brain networks with a stronger left lateralization for the semantic task and a more bilateral pattern for the tone task (Figure S8). When contrasting individual semantic conditions, WPM and FPM, with tone judgment, they each activated a left-lateralized fronto-temporo-occipital network consisting of semantic representation and control regions (Figure 4A&B). Tone judgment, on the other hand, activated a frontoparietal network of cognitive control regions encompassing right middle, superior, and inferior frontal gyri, angular gyrus, and precuneus, confirming the non-verbal, high executive demand of this task (Figure 4A&B). Notably, tone judgment > WPM activated a cluster in the pre-SMA and dorsal anterior cingulate cortex that was not present for tone judgment > FPM, pointing towards the extended cognitive demand for feature-compared with word-picture matching. This was further confirmed by the direct comparison of both semantic conditions showing increased activity for FPM in domain-specific semantic control regions (left pMTG and IFG) and domain-general cognitive control including left SFG and SPL (Figure 4C). Table S5 lists results for all univariate comparisons.

**Figure 4.**
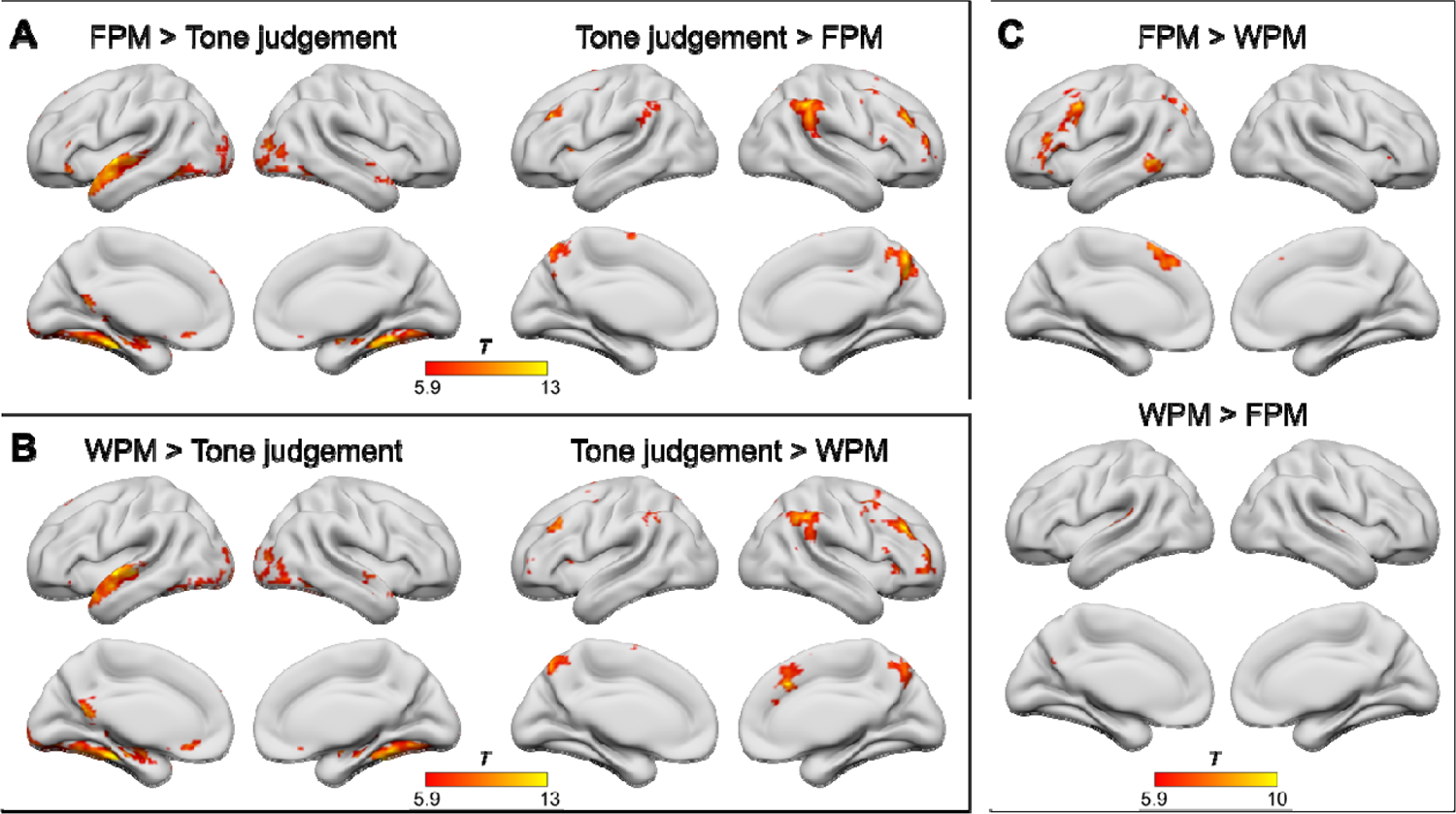
Univariate activation results for experimental conditions during baseline session. (A) Results for feature-picture matching (FPM) contrasted with tone judgment, (B) results for word-picture matching (WPM) contrasted with tone judgment, and (C) contrasting both semantic conditions, WPM and FPM, with each other. Results are FWE-corrected at peak level p < 0.05 with a minimal cluster size k = 10 voxels.

#### iTBS increases task-specific activity for high semantic control demand in the attention network

Results revealed stronger activation only after effective compared with sham sessions and only for both semantic conditions. Compared with rest, there was stronger activation for FPM after effective stimulation in left SPL extending into cuneus and in right AG, occipital, and fusiform gyrus (Figure 5A). When contrasted with tone judgment, a cluster in left occipital lobe and fusiform gyrus was detected (Figure 5A). Table 1 displays detailed results for FPM and WPM. Results for WPM are visualized in Figure S9.

**Figure 5.**
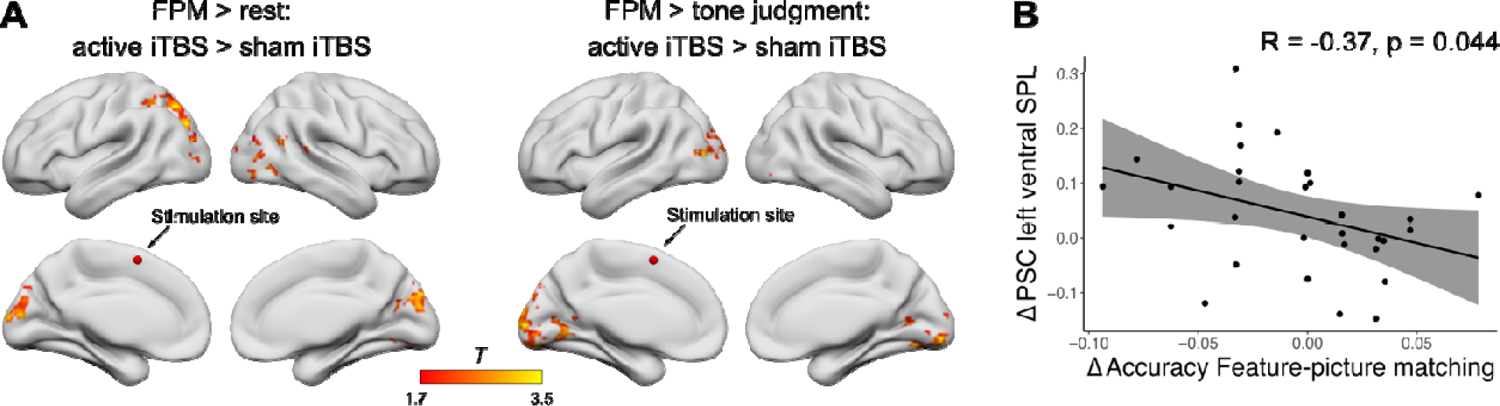
Effect of stimulation on brain activation. After effective stimulation, stronger activation was found for (A) feature-picture matching (FPM) compared with rest (implicit baseline) and tone judgment. We extracted percent signal change (PSC) for significant clusters and correlated the difference in PSC between effective and sham sessions with the difference in behavior between effective and sham sessions. (B) For Δ of accuracy of FPM, a negative correlation with the difference in PSC in the left superior parietal lobe (SPL) was detected. fMRI results are thresholded at p < 0.05 at peak level and FWE-corrected at p < 0.05 at cluster level.

#### Increased activity after iTBS is associated with poorer semantic control

We correlated the difference in PSC (effective > sham iTBS) for the stimulation site pre-SMA and for cluster peaks that showed an effect of stimulation (Table 1) with the difference in behavior. Results showed a negative correlation for accuracy of FPM with a cluster in left dorsal SPL (*r* = −0.37, *p* = 0.044; Figure 5B) such that less PSC after effective iTBS was associated with higher accuracy after iTBS. We did not detect any significant associations with reaction time.

### iTBS increases task-specific connectivity in domain-general default and attention networks

Four seed regions extracted from the univariate stimulation results showed changes in functional connectivity after effective compared with sham stimulation: right cuneus and left ventral SPL, occipital pole, and intra-calcarine cortex. All regions showed greater whole-brain coupling for FPM than WPM after effective stimulation. Additionally, the left SPL also showed greater connectivity for tone judgment > FPM (Figure 6A; Table S6). For the right cuneus, enhanced coupling was found in bilateral STG and frontal opercula, pre-SMA, and right SPL and AG. These clusters were mainly located in dorsal and ventral attention and the right sensorimotor network (Figure 6B). The left occipital pole showed stronger connectivity with clusters in bilateral occipital lobes, which linked to visual networks, and temporal and frontal regions, which were mainly situated in the DMN and sensorimotor networks. The second seed from the occipital cortex, the left intracalcarine cortex, showed enhanced coupling with a cluster in the left MTG and STG, which associated with the DMN. The seed in the left SPL showed different patterns of increased coupling for FPM and tone judgment. For FPM, we found activation in left MTG extending into AG and in the right SPL. Clusters were associated with the temporoparietal subnetwork of the DMN and the parietal control network, respectively.

**Figure 6.**
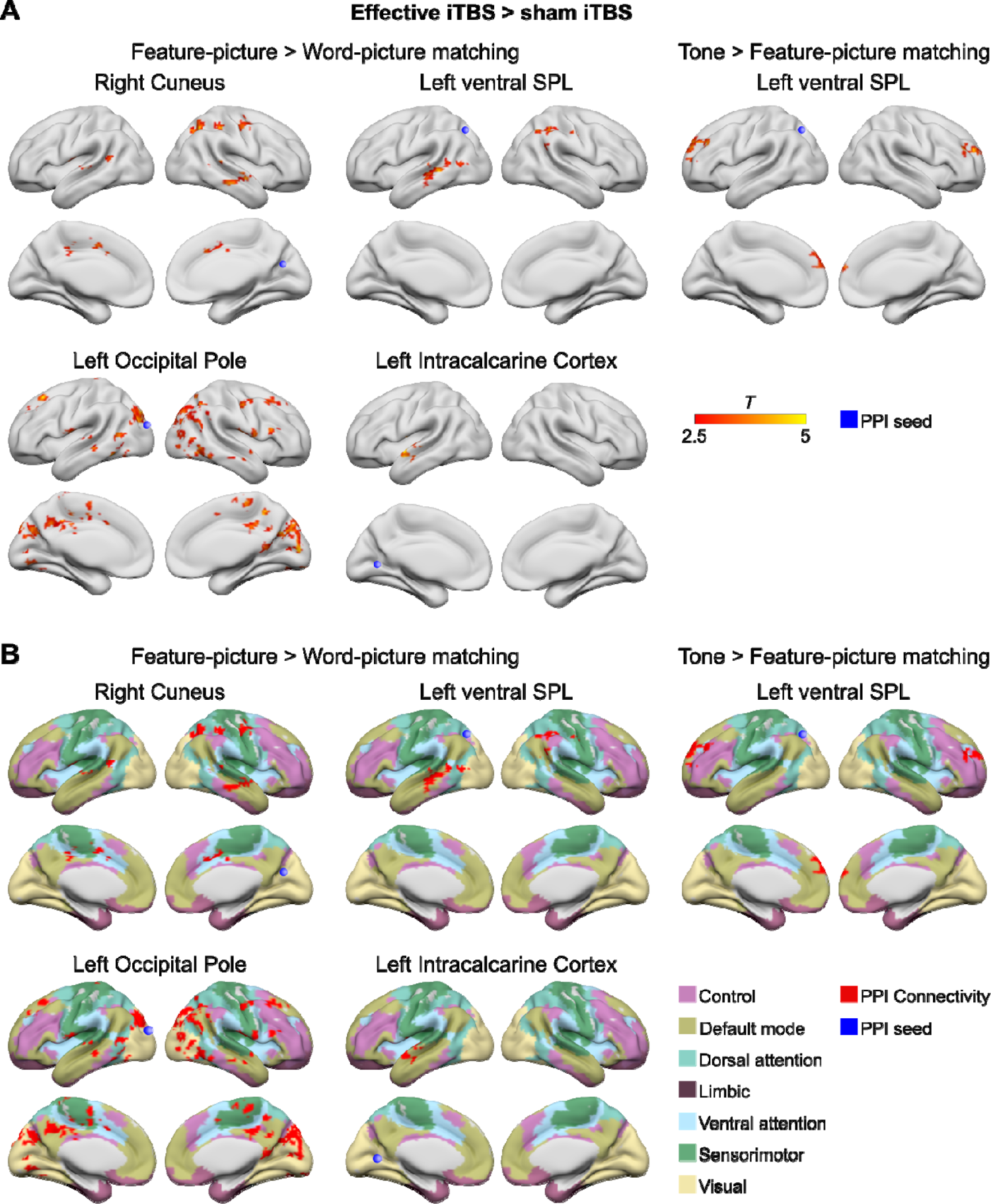
Functional connectivity results for cluster peaks that showed stronger activation after effective iTBS. (A) Four seeds showed increased whole-brain coupling after effective relative to sham stimulation. Only the left ventral SPL seed showed stronger coupling for the task contrasts FPM>WPM and tone judgment>FPM. (B) Binary PPI activation maps plotted onto a seven-networks functional connectivity parcellation (Yeo et al., 2011).

For tone judgment on the other hand, the left SPL coupled with two clusters in the dorsolateral prefrontal cortex, which were both associated with the ventral attention network. Figure 6B shows the overlap of all clusters with a common network parcellation (Yeo et al., 2011).

#### Increased coupling of executive networks after iTBS is associated with faster performance for the most demanding semantic condition

The change in functional connectivity between pre-SMA and left ventral SPL was associated with a change in behavior after effective stimulation (Figure 7). More specifically, a negative correlation (*r* = −0.51, *p* = 0.02 after Bonferroni-Holm correction) indicated that reactions for FPM were slower the more those two regions were decoupled after effective iTBS.

**Figure 7.**
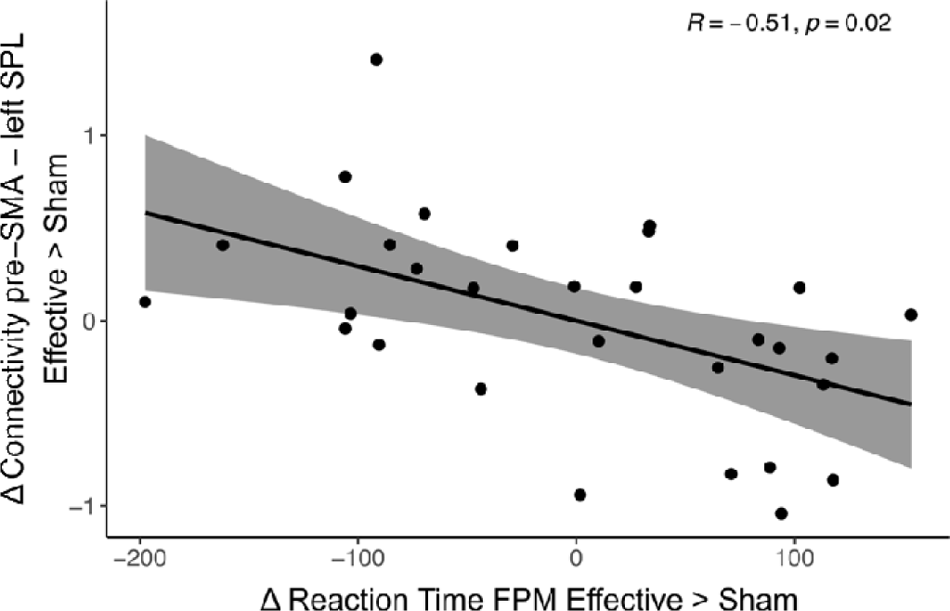
Relationship between stimulation-induced changes in functional connectivity and behavior. Reduced coupling of pre-SMA and left ventral SPL after effective iTBS was associated with slower reaction times (RT) during the feature-picture matching (FPM) condition after effective relative to sham iTBS.

## Discussion

In light of global population aging and the associated increase in age-related diseases, new interventions are needed to counteract cognitive decline and promote successful aging. NIBS is increasingly recognized as a promising tool to boost cognitive functions in older adults. However, to design effective treatment protocols, a better understanding of the neural mechanisms of NIBS is mandatory. In particular, it remains unclear whether stimulation-induced improvements may be underpinned by decreases or increases in task-related activity and connectivity, or both. Here, we explored the effect of effective relative to sham iTBS over the pre-SMA on the behavioral and neural level during a semantic and a tone judgment task. In the absence of direct behavioral changes, we found significant modulation of task-related activity and connectivity. These changes differed in their functional relevance at the behavioral level. Our main results were as follows: iTBS induced increased activation during semantic processing at remote regions in posterior attention and visual networks. Functional connectivity, on the other hand, was modulated by the executive task demand such that high semantic control demands were linked to more widespread coupling with attention, default, and control networks, whereas non-verbal cognitive control led to stronger coupling with attention and control networks. Strikingly, TMS-induced changes on activation and functional connectivity had differential effects on behavior. While increased activity related to poorer semantic control, enhanced coupling between the stimulation site and the dorsal attention network was linked to faster performance in the most demanding semantic condition. Overall, our findings show that iTBS modulates networks in a task-dependent manner and generates effects at regions remote to the stimulation site. Further, our results shed new light on the role of the pre-SMA in domain-general and semantic control processes, indicating that the pre-SMA supports executive aspects of semantic control.

### Higher-order cognitive networks for semantic judgment and tone judgment that overlap in the pre-SMA

Our task paradigm revealed distinct task-specific networks for semantic and tone judgment. Semantic processing activated a left-lateralized fronto-temporal network consisting of semantic representation and control regions [38,39]. Moreover, we found pronounced bilateral activation of middle and posterior fusiform gyri, which have recently been linked to lexical semantics [40]. Contrasting both conditions of semantic judgment with each other validated the intended modulation of cognitive demand: While WPM showed greater activation in left language perception areas, confirming enhanced phonological and lexical processing during this task, FPM activated core regions of semantic but also domain-general control, indicating increased task demand. Contrarily, the tone judgment task activated a frontoparietal network, which strongly overlapped with the MDN, confirming the non-verbal, high executive demand of this task.

### iTBS does not produce direct behavioral changes at the group level

Although univariate fMRI results from the baseline session demonstrated the contribution of the pre-SMA to both experimental tasks, applying effective iTBS to the pre-SMA did not produce direct behavioral changes relative to sham. This result was unexpected. However, we are not the first study to observe stimulation-induced effects on the neural but not behavioral level in healthy older adults [4]. The lack of a behavioral effect might be due to numerous reasons. First, including a separate baseline session may have aggravated the chance of observing a stimulation effect since participants were already familiarized with the paradigm and improved across all conditions after the baseline session. Second, offline stimulation might not have been strong enough to induce behavioral changes. Although we took great care to minimize the time between end of stimulation and begin of task-based fMRI, this might have hindered chances for a behavioral effect. Third, the tasks may have been too easy to observe a facilitatory effect of iTBS. This is the first study to use TBS in healthy aging in semantic cognition. While previous work successfully applied anodal electrical stimulation to the left inferior frontal and motor cortex to enhance semantic processing in older adults [41–43], the facilitatory stimulation of an executive control hub might not have been critical when task performance is already high. Nonetheless, though unintended, the absence of a stimulation effect on cognition allowed us to interpret alterations on the neural level without the confounds of behavioral changes that might make them harder to interpret otherwise [44,45]. Moreover, the behavioral relevance of these changes was demonstrated in the significant correlations between activity or connectivity increases and behavioral modulation.

### iTBS over the pre-SMA increases activity in a widespread network of visual processing and cognitive control

Effective iTBS generated greater activity in posterior regions but not at the stimulation site. This finding was surprising but is in line with previous observations that TBS produces remote effects in neural networks [4,46,47]. Notably, stimulation-induced changes in neural activity were only found for the semantic task and in the occipital cortex, including bilateral lingual gyri and medial occipital lobe. There is emerging evidence from healthy but also patient studies that the occipital cortex, particularly the lingual gyrus, supports language-related and verbal memory tasks [48–51]. Thus, the increased activation of these regions mediated by the pre-SMA suggests a top-down control on visual processing regions in a task-specific manner. Moreover, for the FPM condition, we found additional activation in the dorsal attention network after effective stimulation, which illustrates a functional connection with focused attention, likely semantic-specific [49,52,53].

### Increased functional coupling after iTBS is modulated by task-relevant networks

Functional connectivity analyses showed increased whole-brain coupling for the two conditions with high executive demand, FPM and tone judgment. Notably, which networks showed enhanced coupling was task-dependent. Seeding in a node of the left dorsal attention network, we found increased connectivity with regions in the left temporoparietal subnetwork of the DMN, which is associated with semantic control, and the homologous right dorsal attention network for FPM compared with WPM. For tone judgment compared with FPM, on the other hand, the same seed showed enhanced connectivity with clusters in middle frontal gyrus, which were linked to frontoparietal control and default networks. These findings highlight the potential of iTBS to generate task-specific changes in functional network coupling [4,47,54]. Further, it suggests a stimulation-induced modulation of whole-brain functional connectivity in response to executive and attention demands and supports the notion of the pre-SMA as an organizing hub in the MDN, coordinating the interaction of different cognitive control regions [55].

### iTBS-induced changes in activation and functional connectivity relate differently to behavior

While it might seem surprising that increased activation of the parietal dorsal attention network was linked to poorer accuracy in the semantic task (Figure 8), this finding corroborates the notion that the most efficient task processing is associated with little or no additional functional activation apart from task-specific core regions. This is a common observation in neurocognitive aging, where increased activation and reduced deactivation of domain-general regions have been associated with neural inefficiency, leading to poorer performance across cognitive domains [56,57]. Moreover, better and more efficient behavioral performance due to training-induced activation decreases has been reported in healthy participants [58,59] as well as post-stroke chronic aphasia [60,61]. In our study, task performance was high and remained unchanged after iTBS, indicating a stimulation-induced upregulation of remote cognitive control regions not necessary for efficient task processing.

**Figure 8.**
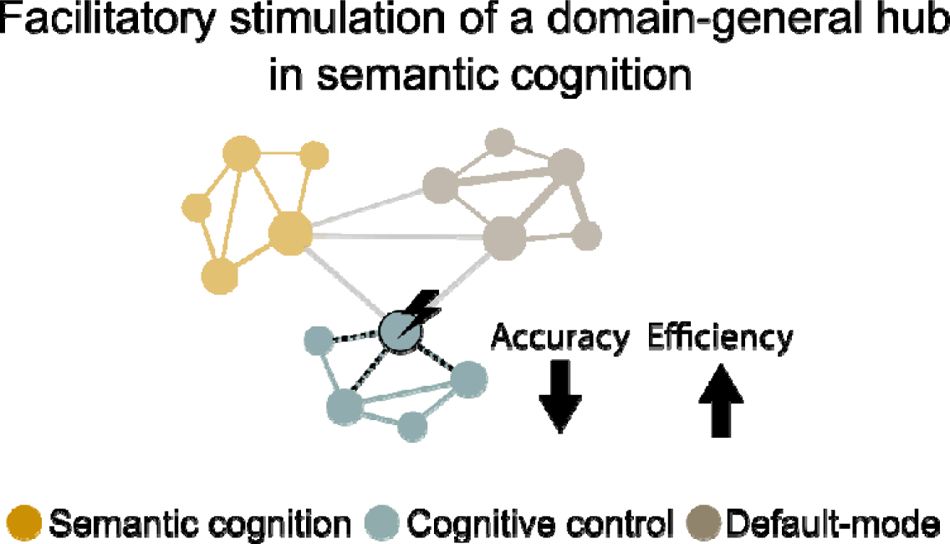
Facilitatory stimulation of a hub of the domain-general multiple-demand network enhanced coupling with other cognitive control networks distal to the stimulation site. Stimulation-induced increases in activity of cognitive control networks were linked to poorer performance, while increased connectivity between task-relevant networks was associated with more efficient processing during semantic judgments.

On the other hand, increasing functional connectivity between our stimulation site and the upregulated cluster in the parietal dorsal attention network after iTBS was associated with faster reactions in the most demanding semantic condition (Figure 8). This result strengthens the idea of a task-specific coupling of cognitive control regions that have been linked to executive components of semantic processing [49,53] and language processing in general [62,63]. Here, we demonstrate that such coupling can enhance the processing efficiency when cognitive demands are high but not the cognitive process per se in form of improved accuracy.

In conclusion, our results agree with the proposal of an adaptive recruitment of domain-general resources to support language processing, which, however, are less efficient than the specialized domain-specific network [2]. Moreover, we demonstrate the potential of facilitating demanding semantic processing in healthy aging via iTBS over a domain-general hub and reveal the underlying modulation of neural networks. Stimulation effects on activity and connectivity were constrained by the cognitive load of a task and took place within specialized networks. This has implications for future studies on the application of rTMS to counteract cognitive decline and highlights the need of a better understanding of the neural network effects of NIBS in general. Stimulation approaches that target functional reorganization within specialized networks as well as remote additional resources might help to design more efficient treatment protocols in the future.

## Declarations of interest

none.

## Acknowledgments

SM held a stipend by the German Academic Scholarship Foundation (Studienstiftung des deutschen Volkes). DS was supported by the Deutsche Forschungsgemeinschaft (SA 1723/5-1) and the James S. McDonnell Foundation (Understanding Human Cognition, #220020292). GH was supported by the Lise Meitner excellence program of the Max Planck Society and the Deutsche Forschungsgemeinschaft (HA 6314/3-1, HA 6314/4-1, HA 6314/4-2). The authors would like to thank the medical technical assistants of MPI CBS for their support with data acquisition.

## Notes

https://identifiers.org/neurovault.collection:13064

### Competing Interest Statement

The authors have declared no competing interest.

### Summary of Updates

Revised manuscript; Supplemental files updated

