## Supplementary materials for "Facilitatory stimulation of the pre-SMA enhances semantic cognition via remote network effects on task-based activity and connectivity"

### Supplementary Methods

#### Description of experimental tasks

#### Semantic judgment task

The semantic judgment task consisted of a word-picture matching (WPM) and a feature-picture matching (FPM) condition, thus varying with respect to the semantic demand of an item. During both conditions, participants listened to a short phrase (e.g., “Is a banana” or “Is sour”) followed by a picture of an object at the offset of the auditory stimulus. They were then asked to judge if the auditory phrase and the presented object match. Stimuli were chosen from eight categories (four living: birds, fruits, mammals, and vegetables; four non-living: clothes, furniture, tools, and vehicles) according to German norm data for semantic typicality (Schröder et al., 2012). From each category, 12 members were selected, 2/3 of them representing typical and 1/3 representing atypical items of the respective category. Hence, in total, 96 stimuli were developed. For each item, a feature from available concept property norms (Devereux et al., 2014) was chosen so that within every category, items could be paired up with regard to their grammatical gender and their feature. In this way, we made sure that every object was introduced with the appropriate gender through the indefinite article both in the congruent and incongruent condition in the auditory stimulus in the WPM condition (“Is a banana” or “Is a lemon”), thus precluding any syntactic clues on accuracy. In German, the indefinite article can take two forms: feminine “eine” and masculine and neutral “ein”. Accordingly, items with male and neutral gender could be paired up together and items with female gender were paired up separately. Further, the arrangement in pairs allowed us to balance the occurrence and to control the semantic value of the features. That is, every feature property was once used as congruent and as incongruent. Since item pairs were within categories, we assured that both congruent and incongruent features were semantically associated with the items. Through this approach, we aimed at avoiding any response bias which could be introduced when an incongruent feature has a bigger semantic distance than the congruent feature from the target item. All auditory stimuli were recorded through the same professional native German female speaker as in the language localizer. Recordings were processed in the same way: They were cut using Praat and normalized via Audacity software.

Across categories, items were balanced for lexical frequency of words and lexical frequency of features using the dlexDB database (Heister et al., 2011). There was no significant difference between frequencies of words ($M$ = 10.27, $SD$ = 23.65) and features ($M$ = 13.63, $SD$ = 23.50), $t(185)$ = 0.98, $p$ = .331). Additionally, items were also balanced across categories for length in phonemes, length in syllables, and length in seconds of the audio files of words and features respectively. In comparison, audio files of words ($M$ = 1.24 s, $SD$

= 0.13) were longer than those of features (*M* = 1.16 s, *SD* = 0.18), $t(176) = 3.81$, $p < .001$). We dealt with this difference in the length of audio files by designing the paradigm in a way that pictures of objects only appeared at the offset of each auditory stimulus, thus not depending the decision-making process on the length of the audio files. Pictures for stimulus items were taken from the freely available Bank of Standardized stimuli (Brodeur et al., 2010, 2014) and the colored picture set by Moreno-Martínez and Montoro (2012) or bought through a license of MPI CBS on Shutterstock. Objects were presented on a white background and all pictures were cropped to a size of 720 x 540 pixels. Stimuli of the semantic judgment task were investigated in a pilot experiment ($n = 50$) beforehand to confirm the intended modulation in task demand and to validate name and feature agreement for each item. Stimuli for the final set were only chosen if they showed at least 80% agreement for the WPM and FPM conditions.

We developed six individual stimuli lists per participant (three sessions with two runs each) such that every item appeared once in every condition across runs and sessions. Conditions and congruency were balanced across runs with pairs of congruent and incongruent stimuli never occurring in the same run. Across participants and runs, accuracy and congruency of individual items were pseudorandomized. After balancing procedures, stimuli lists were randomized.

**Tone judgment task**

The non-verbal control task consisted of sinewave tones at different frequencies (300–825 Hz), which were presented in a sequence of two tones. Tones in a sequence always had a difference in frequency of 250 Hz. Individual tones were generated using a pure tone generator in Matlab with the following parameters: sampling frequency of 16.000 Hz, duration of 450 ms, and fade-in and fade-out duration of 10 ms each. Afterwards, tones were paired up using Audacity software so that each tone once appeared first and once second in a sequence. An inter-tone interval of 300 ms was included in each sequence. Thus, each tone sequence had a length of 1200 ms which equaled the average length of all verbal stimuli. In the control task, participants heard a tone sequence and were asked to match this with a picture of an arrow pointing diagonally upwards or downwards which appeared at the offset of the auditory stimulus. Like in the semantic judgment task, participants had to indicate their choice via button press. Through this process, we aimed at keeping the task as similar as possible to the semantic judgment task but without any verbal processing involved.

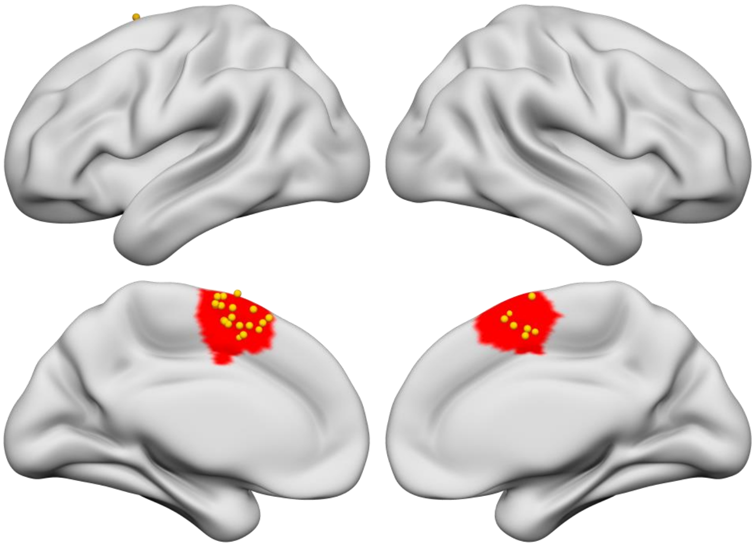

**Figure S1. Individual stimulation sites within a pre-defined mask of the pre-SMA.** The mask was taken from a freely available probabilistic cytoarchitectonic map (Ruan et al., 2018). Yellow nodes signify individual stimulation sites (n = 30). Due to the midline structure of the pre-SMA, we did not restrict sites to one hemisphere within the mask.

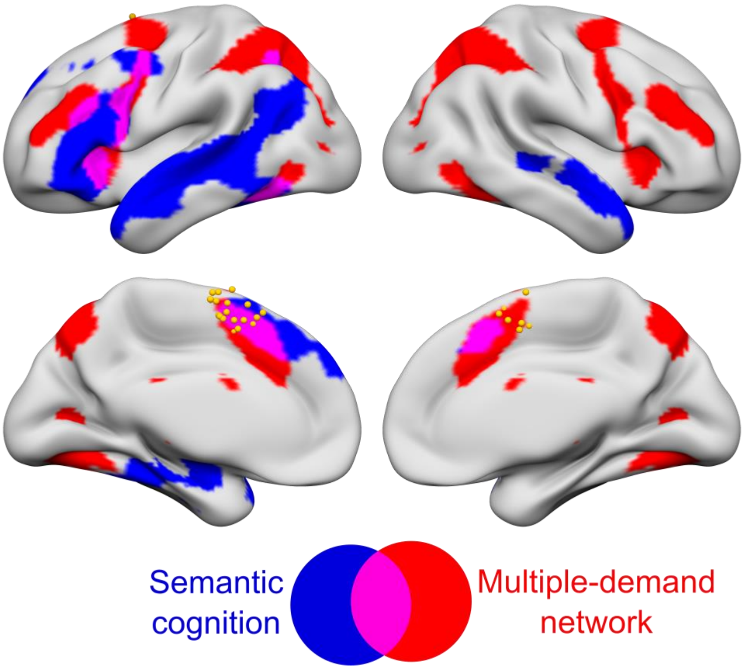

**Figure S2. Individual stimulation sites within the semantic cognition and the multiple-demand network.** The map of the semantic cognition network was taken from Jackson (2021) and the multiple-demand network from Fedorenko et al., 2013.

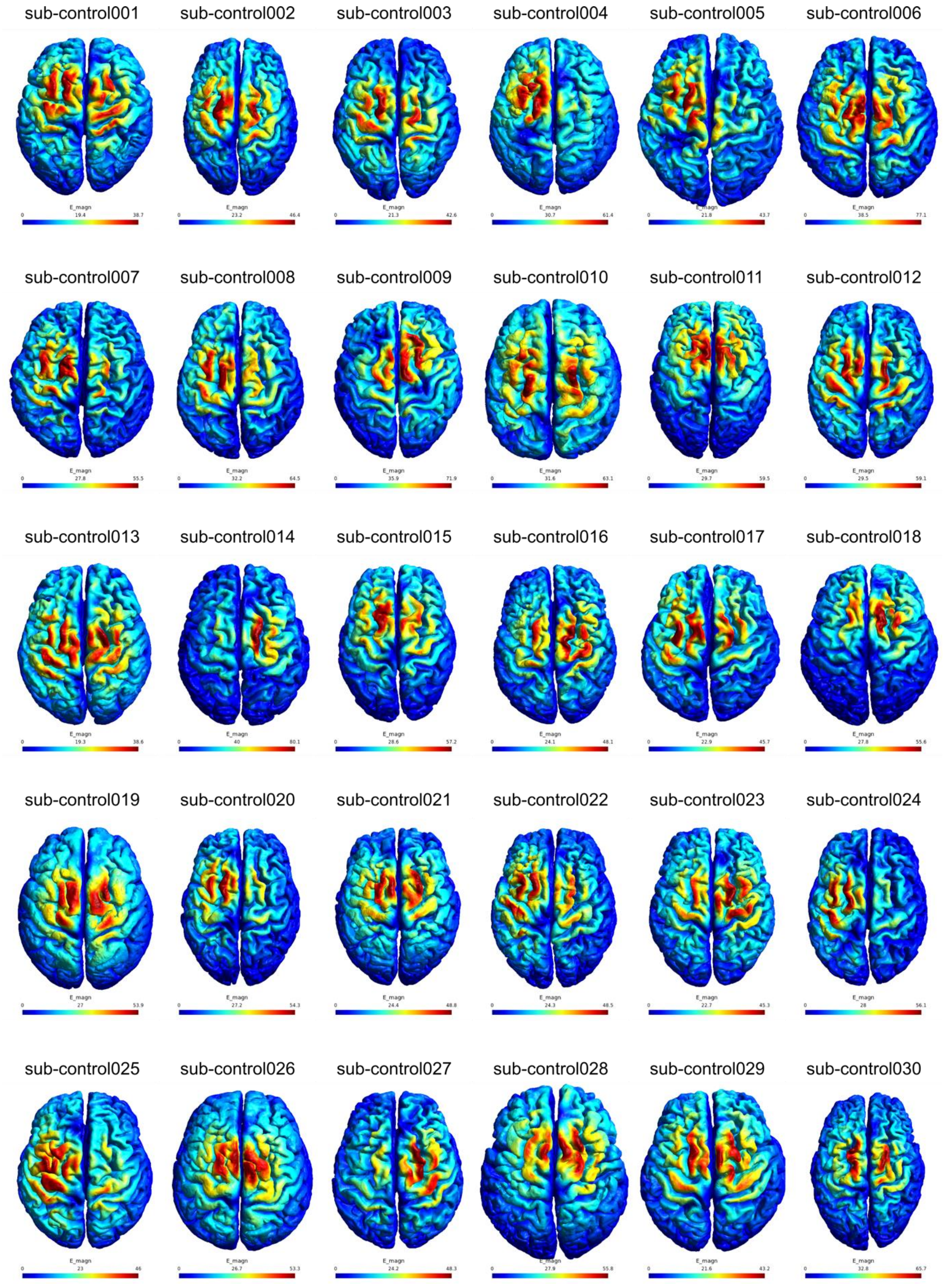

**Figure S3. Results of electrical field simulation.** Individual electrical fields induced by iTBS over the pre-SMA are plotted on the individual surface reconstruction and displayed in axial view.

**Table S1**. Model for DV Accuracy

| Comparison | Model | Δ AIC | df |
|---|---|---|---|
| Random effects | **~ session x condition + (1\|sub)** | **0** | **10** |
| | ~ session x condition | 71.4 | 9 |
| Fixed effects | **~ session x condition x congruency + age + (1\|sub)** | **0** | **20** |
| | ~ session x condition + congruency + (1\|sub) | 55.6 | 11 |
| | ~ session x condition + congruency + age + (1\|sub) | 57.6 | 12 |
| | ~ session x condition + (1\|sub) | 131.6 | 10 |
| | ~ session x condition + task + (1\|sub) | 131.6 | 10 |
| | ~ session x condition + stim_order + (1\|sub) | 133.2 | 11 |
| | ~ session x condition + age + (1\|sub) | 133.6 | 11 |

*Note.* Winning models are highlighted in bold; AIC Akaike Information Criterion

**Table S2.** Model for DV log(RT)

| Comparison | Model | Δ AIC | df |
|---|---|---|---|
| Random effects | **~ session x condition + (1 + session\|sub) + (1\|stimulus_audio)** | 0.0 | 17 |
| | ~ session x condition + (1 + session\|sub) + (1\|stimulus_picture) | 1058.4 | 17 |
| | ~ session x condition + (1 + session\|sub) | 1971.3 | 16 |
| | ~ session x condition + (1\|sub) | 2644.0 | 11 |
| | ~ session x condition | 8421.5 | 10 |
| Fixed effects | **~ session x condition + congruency + age + (1 + session\|sub) + (1\|stimulus_audio)** | **0** | **19** |
| | ~ session x condition + congruency + (1 + session\|sub) + (1\|stimulus_audio)2 | 7.4 | 18 |
| | ~ session x condition + age + (1 + session\|sub) + (1\|stimulus_audio) | 222.8 | 18 |
| | ~ session x condition + (1 + session\|sub) + (1\|stimulus_audio) | 230.3 | 17 |
| | ~ session x condition + task + (1 + session\|sub) + (1\|stimulus_audio) | 230.3 | 17 |
| | ~ session x condition + stim_order + (1 + session\|sub) + (1\|stimulus_audio) | 232.3 | 18 |
| Interactions | **~ session x condition + congruency + condition x congruency + age + (1 + session\|sub) + (1\|stimulus_audio)** | **0** | **21** |
| | ~ session x condition + session x congruency + condition x congruency + age + (1 + session\|sub) + (1\|stimulus_audio) | 2.1 | 23 |
| | ~ session x condition x congruency + age + (1 + session\|sub) + (1\|stimulus_audio) | 8.3 | 27 |

| | | | |
|---|---|---|---|
| | ~ session x condition + congruency + age + (1 + session\|sub) + (1\|stimulus_audio) | 302.5 | 19 |

*Note.* Winning models are highlighted in bold; AIC Akaike Information Criterion

**Preprocessing of MRI data**

Results included in this manuscript come from preprocessing performed using fMRIPrep 20.2.3 (Esteban et al., 2019; Esteban, Blair, et al. (2018); RRID:SCR_016216), which is based on Nipype 1.6.1 (Gorgolewski et al., 2011, 2017).

*Anatomical data preprocessing*

A total of 1 T1-weighted (T1w) images were found within the input BIDS dataset. The T1-weighted (T1w) image was corrected for intensity non-uniformity (INU) with N4BiasFieldCorrection (Tustison et al., 2010), distributed with ANTs 2.3.3 (Avants et al., 2008), and used as T1w-reference throughout the workflow. The T1w-reference was then skull-stripped with a Nipype implementation of the antsBrainExtraction.sh workflow (from ANTs), using OASIS30ANTs as target template. Brain tissue segmentation of cerebrospinal fluid (CSF), white-matter (WM) and gray-matter (GM) was performed on the brain-extracted T1w using fast (FSL 5.0.9, RRID:SCR_002823, Zhang et al., 2001). Brain surfaces were reconstructed using recon-all (FreeSurfer 6.0.1, RRID:SCR_001847, Dale et al., 1999), and the brain mask estimated previously was refined with a custom variation of the method to reconcile ANTs-derived and FreeSurfer-derived segmentations of the cortical gray-matter of Mindboggle (RRID:SCR_002438, Klein et al., 2017). Volume-based spatial normalization to two standard spaces (MNI152NLin6Asym, MNI152NLin2009cAsym) was performed through nonlinear registration with antsRegistration (ANTs 2.3.3), using brain-extracted versions of both T1w reference and the T1w template. The following templates were selected for spatial normalization: FSL's MNI ICBM 152 non-linear 6th Generation Asymmetric Average Brain Stereotaxic Registration Model [Evans et al. (2012), RRID:SCR_002823; TemplateFlow ID: MNI152NLin6Asym], ICBM 152 Nonlinear Asymmetrical template version 2009c [Fonov et al. (2009), RRID:SCR_008796; TemplateFlow ID: MNI152NLin2009cAsym],

*Functional data preprocessing*

For each of the 8 BOLD runs found per subject (across all tasks and sessions), the following preprocessing was performed. First, a reference volume and its skull-stripped version were generated using a custom methodology of fMRIPrep. A B0-nonuniformity map (or fieldmap) was estimated based on two (or more) echo-planar imaging (EPI) references with opposing phase-encoding directions, with 3dQwarp Cox and Hyde (1997) (AFNI 20160207). Based on the estimated susceptibility distortion, a corrected EPI (echo-planar imaging) reference was calculated for a more accurate co-registration with the anatomical reference. The BOLD reference was then co-registered to the T1w reference using bbregister (FreeSurfer) which implements boundary-based registration (Greve and Fischl 2009). Co-registration was configured with six degrees of freedom. Head-motion parameters with respect to the BOLD reference (transformation matrices, and six corresponding rotation and translation parameters) are estimated before any spatiotemporal filtering using mcflirt (FSL 5.0.9, Jenkinson et al. 2002). BOLD runs were slice-time corrected using 3dTshift from AFNI 20160207 (Cox and Hyde 1997, RRID:SCR_005927). The BOLD time-series (including slice-timing correction when applied) were resampled onto their original, native space by applying a single, composite transform to correct for head-motion and susceptibility distortions. These resampled BOLD time-series will be referred to as preprocessed BOLD in original space, or just preprocessed BOLD. The BOLD time-series were resampled into standard space, generating a preprocessed BOLD run in MNI152NLin6Asym space. First, a reference volume and its skull-stripped version were generated using a custom methodology of fMRIPrep. Several confounding time-series were calculated based on the preprocessed BOLD: framewise displacement (FD), DVARS and three region-wise global signals. FD was computed using two formulations following Power (absolute sum of relative motions, Power et al. (2014)) and Jenkinson (relative root mean square displacement between affines, Jenkinson et al. (2002)). FD and DVARS are calculated for each functional run, both using their implementations in Nipype (following the definitions by Power et al. 2014). The three global signals are extracted within the CSF, the WM, and the whole-brain masks. Additionally, a set of physiological regressors were extracted to allow for component-based noise correction (CompCor, Behzadi et al. 2007). Principal components are estimated after high-pass filtering the preprocessed BOLD time-series (using a discrete cosine filter with 128s cut-off) for the two CompCor variants: temporal (tCompCor) and anatomical (aCompCor). tCompCor components are then calculated from the top 2% variable voxels within the brain mask. For aCompCor, three probabilistic masks (CSF, WM and combined CSF+WM) are generated in anatomical space. The implementation differs from that of Behzadi et al. in that instead of eroding the masks by 2 pixels on BOLD space, the aCompCor masks are subtracted a mask of pixels that likely contain a volume fraction of GM. This mask is obtained by dilating a GM mask extracted from the FreeSurfer’s aseg segmentation, and it ensures components are not extracted from voxels containing a minimal fraction of GM. Finally, these masks are resampled into BOLD space and binarized by thresholding at 0.99 (as in the original implementation). Components are also calculated separately within the WM and CSF masks. For each CompCor decomposition, the k components with the largest singular values are retained, such that the retained components’ time series are sufficient to explain 50 percent of variance across the nuisance mask (CSF, WM, combined, or temporal). The remaining components are dropped from consideration. The head-motion estimates calculated in the correction step were also placed within the corresponding confounds file. The confound time series derived from head motion estimates and global signals were expanded with the inclusion of temporal derivatives and quadratic terms for each (Satterthwaite et al. 2013). Frames that exceeded a threshold of 0.5 mm FD or 1.5 standardised DVARS were annotated as motion outliers. All resamplings can be performed with a single interpolation step by composing all the pertinent transformations (i.e. head-motion transform matrices, susceptibility distortion correction when available, and co-registrations to anatomical and output spaces). Gridded (volumetric) resamplings were performed using antsApplyTransforms (ANTs), configured with Lanczos interpolation to minimize the smoothing effects of other kernels (Lanczos 1964). Non-gridded (surface) resamplings were performed using mri_vol2surf (FreeSurfer).

Many internal operations of fMRIPrep use Nilearn 0.6.2 (Abraham et al. 2014, RRID:SCR_001362), mostly within the functional processing workflow. For more details of the pipeline, see the section corresponding to workflows in fMRIPrep’s documentation.

**Analysis of subject-specific functional regions of interest**

The definition of fROIs followed the procedure described by (Fedorenko et al., 2010) and was done using the spm_ss toolbox (Nieto-Castañón & Fedorenko, 2012): First, individual activation maps for our contrast of interest of the localizer task (intact > acoustically degraded speech) were thresholded at a voxel-wise false discovery rate (FDR) of $q < 0.05$ at whole-brain level (Genovese et al., 2002) and then overlaid on top of each other. The resulting probabilistic overlap map displayed how many participants showed activation at each voxel. Next, the overlap map was smoothed (5 mm), thresholded at 3 participants (10%; cf. Fedorenko et al., 2010), and parcellated using a watershed algorithm (Meyer, 1991). The watershed algorithm resulted in 37 fROIs. Third, only those ROIs from the parcellation were retained where at least 60% of participants had any supra-threshold voxels (cf. Fedorenko et al., 2010; Julian et al., 2012). This led to a final sample of 25 parcels (Figure S4). To confirm that these parcels were indeed relevant to language processing, independent of the task, we entered them in a random-effects group-level analysis using the experimental task data. Results were calculated for the contrast language (i.e., WPM + FPM) > rest and FDR-corrected at $q < 0.05$. Results showed that all 25 parcels were significantly stronger activated for the language task. Finally, subject-specific fROIs were defined as the 10% most active voxels in each participant for the localizer contrast intact > degraded speech within each parcel. Since we were interested in the potential effect of iTBS on differences in activation in the language-specific fROIs, we extracted PSC for each fROI and condition for the experimental task using the MarsBar toolbox (version 0.45; Brett et al., 2002). The data were then entered into a linear model with predictors for stimulation type (effective or sham) and fROI and their interaction term. Post-hoc comparisons were applied using the package emmeans (Lenth, 2020).

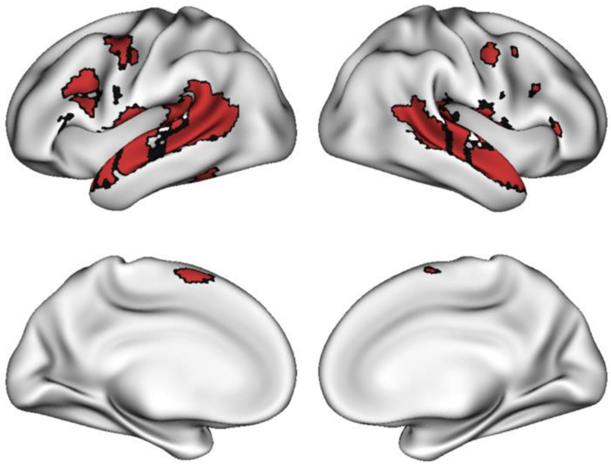

**Figure S4. Results of group-constrained subject-specific parcellation approach for language localizer task.** The figure shows 25 regions of interest (ROIs) that were activated in at least 60% of participants for the contrast intact > degraded speech after data were parcellated using a watershed algorithm. To confirm that these parcels were indeed relevant to language processing, independent of the task, we entered them in a random-effects group-level analysis using the independent data set for the semantic judgement task at baseline. Results showed that all 25 parcels were significantly stronger activated for the language task relative to rest (see Table S3).

**Table S3.** Results of random-effects analysis semantic judgement > rest for 25 functional ROIs from localizer contrast

| ROI | Hemi | Name | Average ROI size | Avg loc. mask size | Overlap | *T* | *Df* | *P* |
|---|---|---|---|---|---|---|---|---|
| 1 | R | pMTG | 473 | 132 | 1.00 | 11.21 | 27.08 | < 0.001 |
| 2 | R | MTG/STG | 334 | 100 | 0.97 | 12.61 | 26.90 | < 0.001 |
| 3 | R | aMTG/ATL | 705 | 195 | 1.00 | 13.05 | 27.20 | < 0.001 |
| 4 | L | MTG | 581 | 157 | 1.00 | 13.40 | 28.35 | < 0.001 |
| 5 | L | ATL | 204 | 52 | 0.97 | 13.35 | 26.86 | < 0.001 |
| 6 | R | Operculum | 168 | 42 | 0.93 | 10.31 | 25.36 | < 0.001 |
| 7 | L | pMTG/AG/SMG | 1045 | 258 | 1.00 | 13.49 | 27.97 | < 0.001 |
| 8 | L | pMTG/STG | 86 | 24 | 0.97 | 12.81 | 26.63 | < 0.001 |
| 9 | L | Temporal pole | 126 | 32 | 0.97 | 11.72 | 26.00 | < 0.001 |
| 10 | L | STG | 16 | 5 | 0.93 | 15.05 | 25.36 | < 0.001 |
| 11 | L | pMTG | 19 | 6 | 0.93 | 9.67 | 26.27 | < 0.001 |
| 12 | L | pMTG | 14 | 4 | 0.80 | 6.51 | 22.38 | < 0.001 |
| 13 | L | Operculum | 210 | 41 | 1.00 | 11.95 | 25.44 | < 0.001 |
| 14 | R | Operculum | 252 | 52 | 0.87 | 12.32 | 21.76 | < 0.001 |
| 15 | L | Precentr. Gyrus | 263 | 52 | 1.00 | 12.37 | 26.24 | < 0.001 |
| 16 | R | Precentr. gyrus | 142 | 29 | 0.73 | 9.27 | 19.72 | < 0.001 |
| 17 | R | AG | 97 | 18 | 0.77 | 2.57 | 20.11 | 0.00914 |
| 18 | L | IFG, pars op. | 147 | 25 | 0.90 | 12.58 | 22.40 | < 0.001 |
| 19 | R | Cerebellum | 113 | 21 | 0.70 | 8.81 | 18.79 | < 0.001 |
| 20 | L | IFG, pars op. | 79 | 17 | 0.63 | 10.61 | 16.42 | < 0.001 |
| 21 | R | Cerebellum | 129 | 22 | 0.90 | 9.13 | 23.09 | < 0.001 |
| 22 | L | Pre-SMA | 73 | 14 | 0.77 | 11.87 | 18.97 | < 0.001 |
| 24 | R | IFG, pars tr. | 130 | 23 | 0.87 | 7.31 | 21.50 | < 0.001 |
| 25 | L | IFG, pars tr. | 59 | 11 | 0.70 | 8.97 | 18.18 | < 0.001 |
| 29 | L | IFG, pars tr. | 35 | 6 | 0.70 | 6.27 | 18.02 | < 0.001 |

*Note:* Df Degrees of freedom; p-value after FDR-correction q < 0.05.

**The effect of iTBS on subject-specific functional ROIs for language processing**

We extracted PSC for effective and sham iTBS sessions in the 25 subject-specific functional ROIs that were defined using a group-constrained subject-specific parcellation approach. We were interested in an effect of iTBS on PSC of the different conditions. To this end, we fitted linear mixed-effects models with predictors for session and PSC. We did not find any significant interaction between functional ROIs and session. Figures S5-7 show the individual PSC for both stimulation sessions for each ROI and condition.

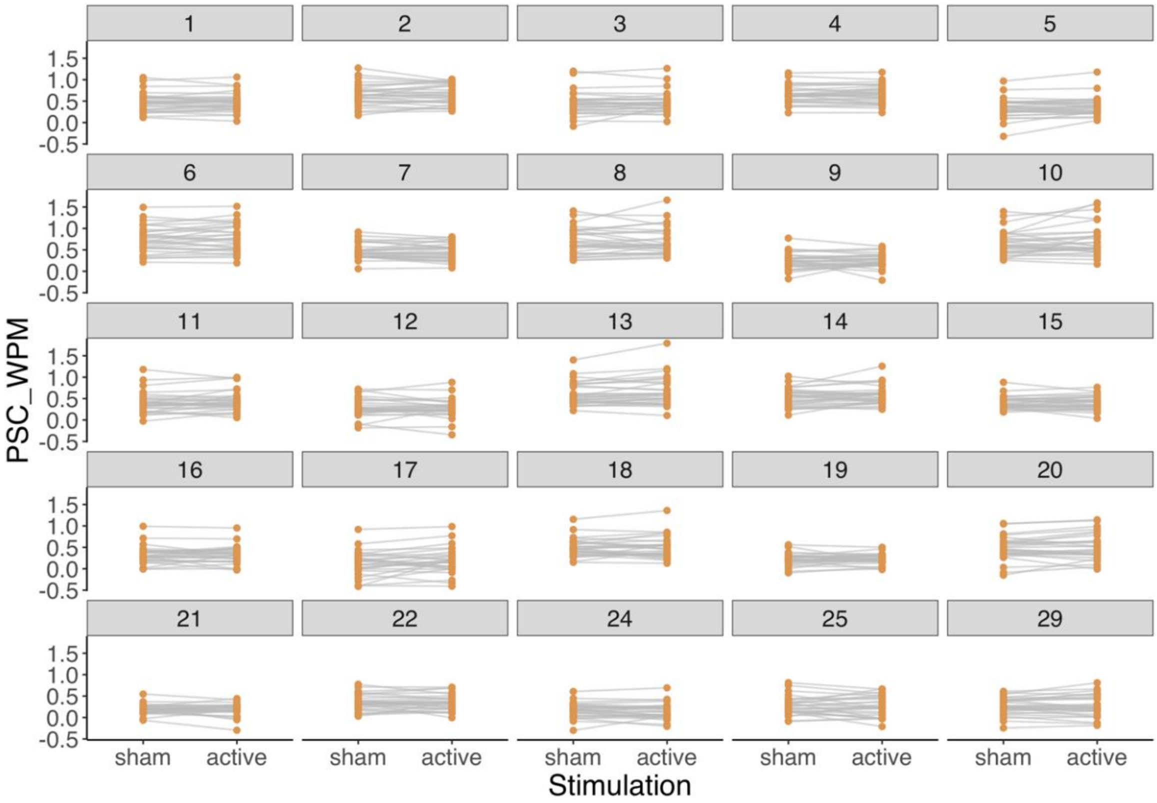

**Figure S5.** PSC in word-picture matching in 25 regions of interest according to subject-specific group parcellation for language localizer task.

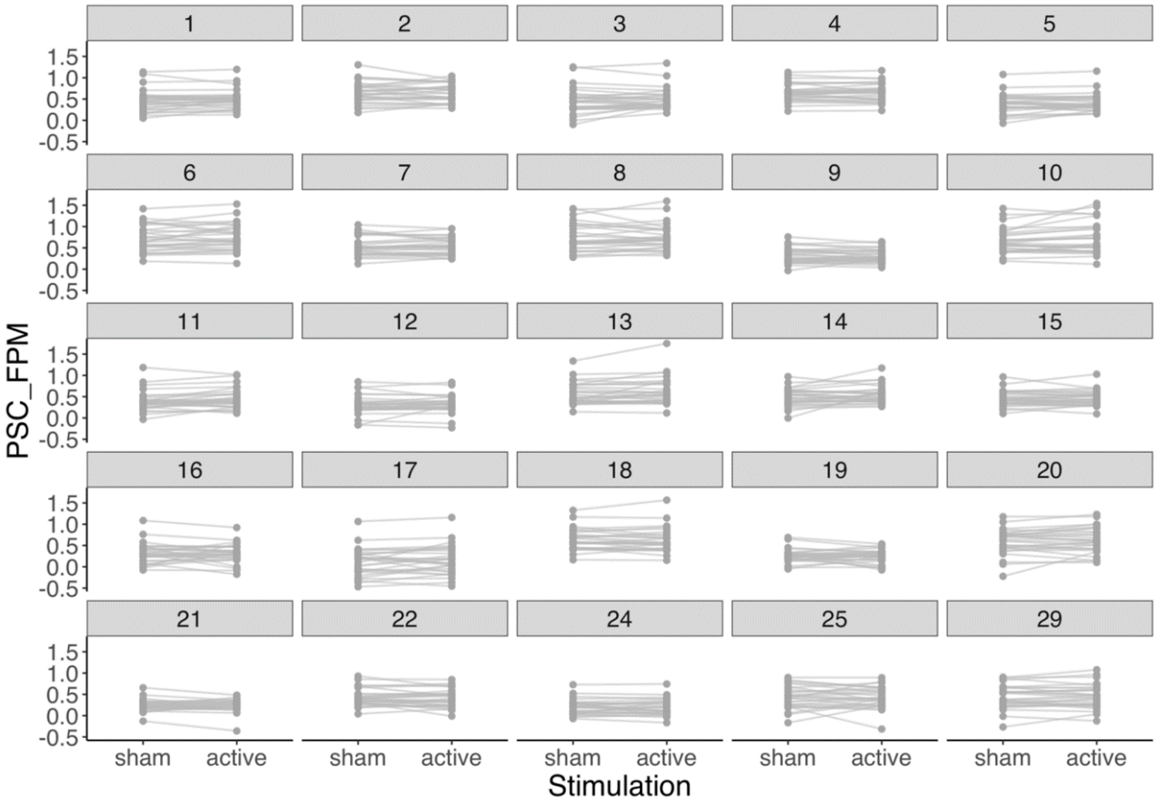

**Figure S6.** PSC in word-picture matching in 25 regions of interest according to subject-specific group parcellation for language localizer task.

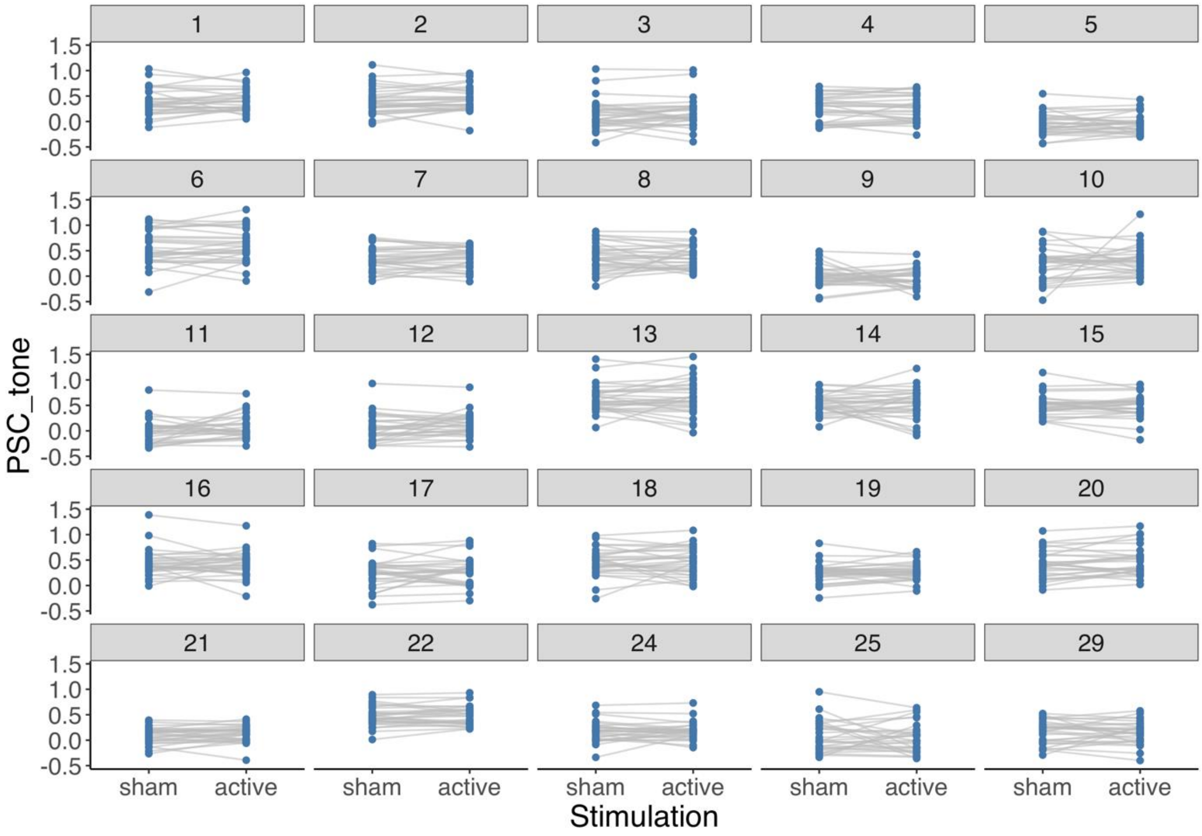

**Figure S7.** PSC in word-picture matching in 25 regions of interest according to subject-specific group parcellation for language localizer task.

### Supplementary Results

**Table S4.** Results of mixed-effects models for accuracy and reaction time

| | Accuracy | | | | Reaction time | | | |
|---|---|---|---|---|---|---|---|---|
| *Coefficient* | *Log-Odds* | *CI* | *Statistic* | *p* | *Estimates* | *CI* | *Statistic* | *p* |
| Intercept | 3.44 | 3.25 – 3.62 | 36.71 | **<0.001** | 6.90 | 6.85 – 6.96 | 242.35 | **<0.001** |
| Session: baseline | -0.62 | -1.03 – -0.21 | -2.95 | **0.003** | 0.19 | 0.11 – 0.27 | 4.75 | **<0.001** |
| Session: active | 0.34 | -0.10 – 0.78 | 1.52 | 0.128 | -0.09 | -0.13 – -0.04 | -3.75 | **<0.001** |
| Condition: WPM | 3.04 | 2.51 – 3.57 | 11.22 | **<0.001** | -0.30 | -0.37 – -0.23 | -8.54 | **<0.001** |
| Condition: FPM | -2.12 | -2.49 – -1.76 | -11.33 | **<0.001** | 0.39 | 0.32 – 0.45 | 11.98 | **<0.001** |
| Congruency: congruent | -0.64 | -0.85 – -0.43 | -6.05 | **<0.001** | -0.04 | -0.06 – -0.03 | -6.36 | **<0.001** |
| Age | 0.00 | -0.02 – 0.02 | 0.07 | 0.943 | | | | |
| Condition WPM * Congruency congruent | 0.63 | -1.48 – 2.73 | 0.58 | 0.561 | -0.14 | -0.20 – -0.08 | -4.48 | **<0.001** |
| Condition FPM * Congruency congruent | -1.48 | -3.67 – 0.72 | -1.32 | 0.187 | 0.04 | -0.02 – 0.10 | 1.30 | 0.195 |
| Session baseline * Condition WPM | 2.04 | 0.57 – 3.52 | 2.71 | **0.007** | -0.08 | -0.14 – -0.02 | -2.51 | **0.012** |
| Session active * Condition WPM | -1.22 | -2.78 – 0.34 | -1.54 | 0.124 | 0.06 | -0.00 – 0.12 | 1.92 | 0.055 |
| Session baseline * Condition FPM | 0.70 | -0.12 – 1.52 | 1.67 | 0.095 | | | | |
| Session active * Condition FPM | -0.36 | -1.24 – 0.52 | -0.81 | 0.417 | | | | |
| Age | -1.08 | -2.14 – -0.02 | -1.99 | **0.046** | 0.10 | 0.06 – 0.14 | 4.51 | **<0.001** |
| Session: baseline * Congruency congruent | -1.59 | -2.33 – -0.86 | -4.25 | **<0.001** | 0.24 | 0.17 – 0.31 | 6.54 | **<0.001** |

| | | | | | | | | |
|---|---|---|---|---|---|---|---|---|
| Session: active * Congruency congruent | 0.99 | -3.20 – 5.18 | 0.46 | 0.643 | | | | |
| Session: baseline * Condition: WPM * Congruency congruent | 1.69 | -2.64 – 6.01 | 0.76 | 0.444 | | | | |
| Session: active * Condition: WPM * Congruency congruent | 2.40 | -0.54 – 5.35 | 1.60 | 0.110 | | | | |
| Session: baseline * Condition: FPM * Congruency congruent | -2.22 | -5.32 – 0.87 | -1.41 | 0.159 | | | | |
| Session: active * Condition: WPM * Congruency congruent | | | | | 0.09 | 0.04 – 0.14 | 3.40 | **0.001** |
| **Random Effects** | | | | | | | | |
| $\sigma^2$ | 3.29 | | | | 0.05 | | | |
| $\tau_{00}$ | 0.17 sub | | | | 0.01 stimulus_audio | | | |
| | | | | | 0.02 sub | | | |
| $\tau_{11}$ | | | | | 0.05 sub.session1 | | | |
| | | | | | 0.01 sub.session2 | | | |
| $\rho_{01}$ | | | | | -0.31 sub.session1 | | | |
| | | | | | 0.30 sub.session2 | | | |
| ICC | 0.05 | | | | 0.45 | | | |
| Observations | 15750 | | | | 15028 | | | |
| Marginal $R^2$ / Conditional $R^2$ | 0.201 / 0.240 | | | | 0.198 / 0.560 | | | |

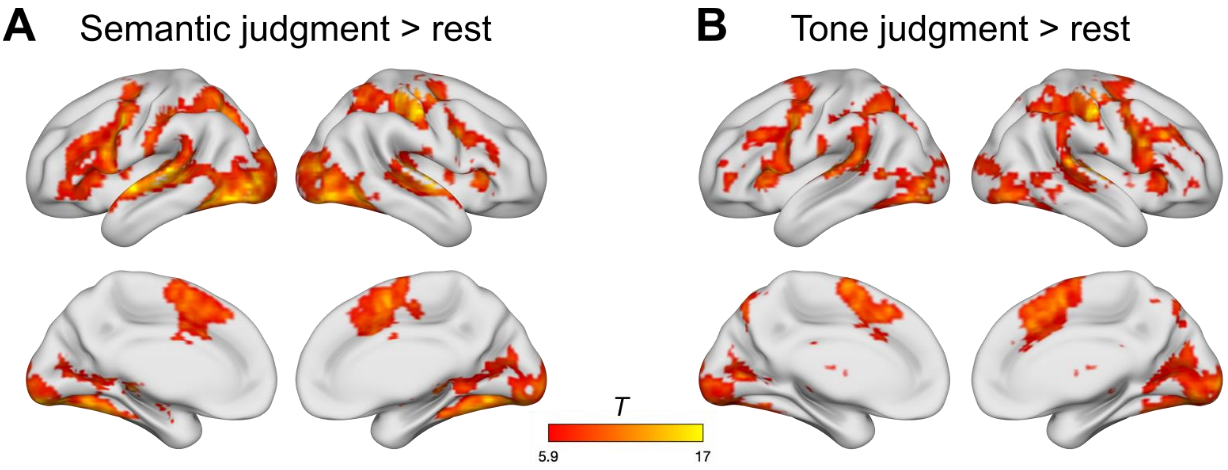

**Figure S8.** Univariate activation results for experimental tasks during baseline session. (A) Brain network for semantic judgment compared with rest (implicit baseline), and (B) brain network for tone judgment compared with rest. Results are FWE-corrected at peak level $p < 0.05$ with a minimal cluster size $k = 10$ voxels.

**Table S5.** Univariate fMRI results – Baseline session

| Anatomical structure | Hemi | *k* | *t* | *x* | *y* | *z* |
|---|---|---|---|---|---|---|
| **Semantic judgment (WPM & FPM) > Rest** | | | | | | |
| Superior temporal gyrus | L | 19000 | 17.45 | -57.66 | -1.60 | -3.25 |
| Cerebellum | L | | 17.41 | -20.34 | -51.36 | -19.75 |
| Postcentral gyrus | R | | 15.33 | 51.82 | -19.02 | 46.25 |
| Supplementary motor area | R | 1074 | 13.25 | 4.54 | 3.38 | 57.25 |
| Presupplementary motor area | R | | 12.00 | 4.54 | 10.84 | 43.50 |
| Supplementary motor area | L | | 11.69 | -7.90 | -1.60 | 60.00 |
| **Tone judgment > Rest** | | | | | | |
| Postcentral gyrus | R | 17880 | 17.61 | 49.33 | -21.50 | 46.25 |
| Superior temporal gyrus | R | | 15.52 | 51.82 | -21.50 | 7.75 |
| Cerebellum | L | | 15.46 | -17.85 | -51.36 | -19.75 |
| Thalamus | L | 13 | 10.55 | -2.92 | -26.48 | -3.25 |
| Precuneus | R | 106 | 9.08 | 14.50 | -66.29 | 46.25 |
| Precuneus | R | | 6.91 | 14.50 | -73.75 | 38.00 |
| Frontal pole | L | 129 | 9.03 | -47.70 | 45.67 | 5.00 |
| Middle frontal gyrus | L | | 8.84 | -37.75 | 43.18 | -0.50 |
| Frontal pole | L | | 8.20 | -42.73 | 55.62 | 2.25 |
| Precuneus | R | 11 | 7.10 | 4.54 | -56.34 | 54.50 |
| Thalamus | L | 11 | 7.06 | -12.87 | -26.48 | -6.00 |
| **WPM > FPM** | | | | | | |
| Superior temporal gyrus | L | 56 | 7.63 | -50.19 | -19.02 | 5.00 |
| Planum temporale | L | | 7.51 | -55.17 | -33.94 | 13.25 |
| Superior temporal gyrus | L | | 6.16 | -45.22 | -26.48 | 5.00 |
| Planum temporale | R | 12 | 7.04 | 54.30 | -21.50 | 10.50 |
| Superior temporal gyrus | R | | 6.26 | 44.35 | -26.48 | 10.50 |
| **FPM > WPM** | | | | | | |
| Middle frontal gyrus | L | 483 | 10.48 | -45.22 | 10.84 | 40.75 |
| Inferior frontal gyrus, pars triangularis | L | | 10.24 | -52.68 | 20.79 | 24.25 |
| Inferior frontal gyrus, pars triangularis | L | | 9.09 | -52.68 | 38.21 | 5.00 |
| Middle temporal gyrus | L | 166 | 9.73 | -62.63 | -51.36 | -6.00 |
| Middle temporal gyrus | L | | 8.56 | -65.12 | -46.38 | 2.25 |
| Inferior temporal gyrus | L | | 8.41 | -52.68 | -58.82 | -8.75 |

| Anatomical structure | Hemi | *k* | *t* | *x* | *y* | *z* |
|---|---|---|---|---|---|---|
| Inferior parietal lobe | L | 156 | 9.45 | -30.29 | -78.73 | 43.50 |
| Middle occipital gyrus | L | | 8.25 | -27.80 | -71.26 | 32.50 |
| Superior parietal lobe | L | | 8.09 | -30.29 | -66.29 | 49.00 |
| Supplementary motor area | L | 122 | 7.96 | -5.41 | 15.82 | 60.00 |
| Superior frontal gyrus | L | | 7.75 | -5.41 | 33.23 | 46.25 |
| Cerebellum | R | 10 | 7.93 | 36.89 | -66.29 | -28.00 |
| Middle frontal gyrus | L | 42 | 7.87 | -35.26 | 8.35 | 60.00 |
| Superior frontal gyrus | L | | 7.73 | -27.80 | 15.82 | 60.00 |
| Middle frontal gyrus | L | | 6.49 | -27.80 | 13.33 | 49.00 |
| **FPM > Tone judgment** | | | | | | |
| Superior temporal gyurs | L | 531 | 13.63 | -57.66 | -1.60 | -6.00 |
| Temporal pole | L | | 11.56 | -52.68 | 13.33 | -17.00 |
| Middle temporal gyrus | L | | 10.76 | -62.63 | -16.53 | -3.25 |
| Fusiform gyrus | L | 900 | 13.15 | -37.75 | -33.94 | -22.50 |
| Fusiform gyrus | L | | 12.03 | -30.29 | -36.43 | -19.75 |
| Fusiform gyrus | L | | 10.07 | -32.78 | -43.90 | -17.00 |
| Fusiform gyrus | R | 677 | 12.21 | 34.40 | -38.92 | -22.50 |
| Fusiform gyrus | R | | 11.49 | 29.42 | -46.38 | -17.00 |
| Parahippocampal cortex | R | | 10.45 | 29.42 | -31.46 | -19.75 |
| Amygdala | R | 21 | 10.31 | 26.94 | -4.09 | -17.00 |
| Precuneus | L | 62 | 10.29 | -5.41 | -56.34 | 16.00 |
| Orbital cortex | L | 41 | 10.28 | -37.75 | 35.72 | -11.50 |
| Inferior frontal gyrus, pars triangularis | L | | 6.19 | -47.70 | 28.26 | -3.25 |
| Temporal pole | R | 148 | 9.24 | 56.79 | 5.86 | -8.75 |
| Temporal pole | R | | 8.67 | 54.30 | 8.35 | -19.75 |
| Temporal pole | R | | 8.15 | 51.82 | 15.82 | -17.00 |
| Frontal pole | L | 59 | 8.13 | -7.90 | 60.60 | 32.50 |
| Frontal pole | L | | 7.30 | -7.90 | 53.14 | 43.50 |
| Frontal pole | L | | 7.21 | -12.87 | 45.67 | 46.25 |
| Pre-SMA | L | 34 | 7.73 | -2.92 | 30.74 | -14.25 |
| Pre-SMA | L | | 6.70 | -0.43 | 40.70 | -11.50 |
| Posterior cingulate cortex | L | 20 | 7.56 | -17.85 | -36.43 | 2.25 |
| Thalamus | L | | 6.50 | -10.38 | -33.94 | 5.00 |
| Inferior frontal gyrus, pars triangularis | L | 21 | 7.19 | -50.19 | 30.74 | 10.50 |

| Anatomical structure | Hemi | *k* | *t* | *x* | *y* | *z* |
|---|---|---|---|---|---|---|
| **Tone judgment > FPM** | | | | | | |
| Precuneus | R | 574.00 | 13.03 | 12.01 | -66.29 | 49.00 |
| Precuneus | R | | 12.49 | 4.54 | -71.26 | 49.00 |
| Precuneus | R | | 12.28 | 7.03 | -76.24 | 40.75 |
| Middle frontal gyrus | L | 81.00 | 12.30 | -37.75 | 33.23 | 32.50 |
| Middle frontal gyrus | L | | 8.58 | -40.24 | 25.77 | 38.00 |
| Frontal pole | R | 300.00 | 11.84 | 39.38 | 38.21 | 32.50 |
| Frontal pole | R | | 10.79 | 41.86 | 43.18 | 24.25 |
| Frontal pole | R | | 9.80 | 39.38 | 53.14 | 13.25 |
| Angular gyrus | R | 661.00 | 10.99 | 54.30 | -46.38 | 40.75 |
| Angular gyrus | R | | 9.20 | 41.86 | -51.36 | 46.25 |
| Angular gyrus | R | | 9.17 | 49.33 | -46.38 | 24.25 |
| Middle frontal gyrus | R | 149.00 | 9.45 | 34.40 | 8.35 | 57.25 |
| Precentral gyrus | R | | 7.39 | 51.82 | -1.60 | 46.25 |
| Precentral gyrus | R | | 7.34 | 46.84 | 3.38 | 51.75 |
| Insula | L | 43.00 | 9.18 | -30.29 | 15.82 | 7.75 |
| Cerebellum | L | 48.00 | 8.71 | -35.26 | -63.80 | -30.75 |
| Cerebellum | L | | 7.03 | -25.31 | -66.29 | -28.00 |
| Inferior frontal gyrus, pars opercularis | R | 27.00 | 8.60 | 56.79 | 15.82 | 5.00 |
| Inferior frontal gyrus, pars opercularis | R | | 6.91 | 51.82 | 15.82 | 13.25 |
| Supramarginal gyrus | L | 100.00 | 8.10 | -50.19 | -51.36 | 38.00 |
| Supramarginal gyrus | L | | 7.53 | -60.14 | -43.90 | 35.25 |
| Supramarginal gyrus | L | | 7.16 | -52.68 | -43.90 | 40.75 |
| Pre-SMA | L | 34.00 | 7.77 | -7.90 | -1.60 | 65.50 |
| Cerebellum | L | 10.00 | 7.23 | -35.26 | -48.87 | -44.50 |
| Middle frontal gyrus | L | 17.00 | 7.13 | -30.29 | -1.60 | 57.25 |
| Cerebellum | L | 17.00 | 6.64 | -35.26 | -63.80 | -44.50 |
| Superior frontal gyrus | R | 10.00 | 6.23 | 16.98 | 3.38 | 65.50 |
| Superior frontal gyrus | R | | 6.21 | 14.50 | -4.09 | 73.75 |

*Note.* Results are FWE-corrected at peak-level at $p < 0.05$. WPM: word-picture matching; FPM: feature-picture matching.

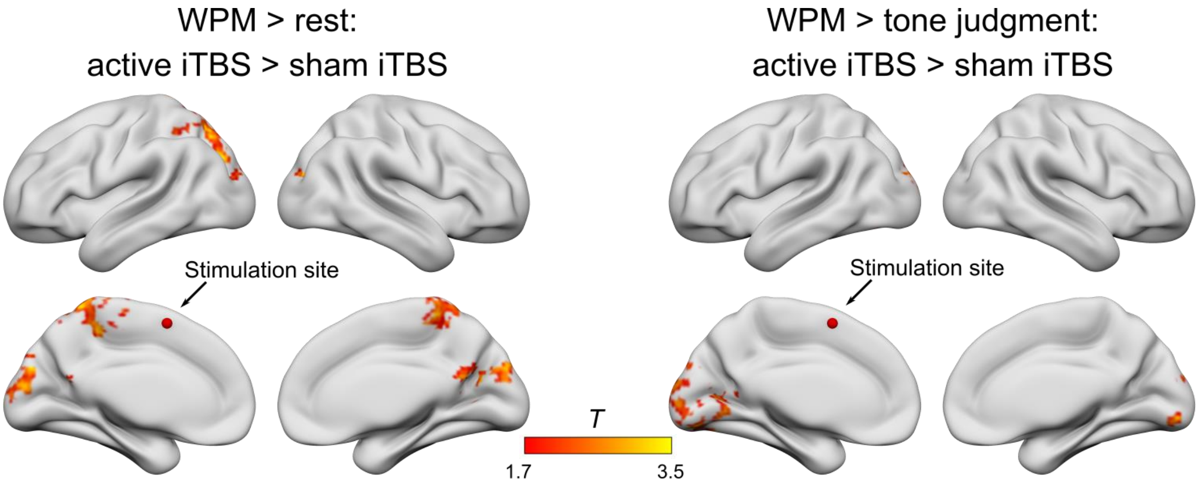

**Figure S9**. Effect of stimulation on brain activation for word-picture matching condition (WPM). After effective stimulation, stronger activation was found for WPM compared with rest and with tone judgment. The activated networks were similar to results for the more demanding feature-picture matching (see Figure 5 in manuscript), except for an additional cluster in the cuneus bilaterally associating with dorsal and ventral attention networks.

**Table S6.** Significant clusters – gPPI Effective > sham iTBS

| **Anatomical structure** | **Hemi** | ***k*** | ***t*** | ***x*** | ***y*** | ***z*** |
|---|---|---|---|---|---|---|
| **Seed: Right Cuneus [9.5, -68.8, 21.5]** | | | | | | |
| **FPM > WPM** | | | | | | |
| Operculum | L | 277 | 5.81 | -57.66 | -9.06 | 7.75 |
| Supramarginal gyrus | L | | 5.02 | -62.63 | -43.90 | 7.75 |
| Superior temporal gyrus | L | | 4.23 | -65.12 | -14.04 | 2.25 |
| Supramarginal gyrus | L | | 3.58 | -50.19 | -48.87 | 16 |
| Superior temporal gyrus | L | | 3.56 | -62.63 | -23.99 | 2.25 |
| Pre-SMA | L | 265 | 5.33 | -0.43 | -6.58 | 46.25 |
| Pre-SMA | R | | 4.80 | 12.01 | 18.30 | 38 |
| Pre-SMA | R | | 4.73 | 12.01 | 8.35 | 38 |
| Anterior cingulate cortex | L | | 3.83 | -7.90 | -23.99 | 43.5 |
| Anterior cingulate cortex | L | | 3.79 | -2.92 | 15.82 | 38 |
| Angular gyrus | R | 197 | 4.87 | 29.42 | -61.31 | 46.25 |
| Superior parietal lobe | R | | 4.41 | 39.38 | -36.43 | 49 |
| Lateral occipital cortex superior | R | | 3.91 | 24.45 | -68.78 | 46.25 |
| Lateral occipital cortex superior | R | | 3.81 | 24.45 | -61.31 | 54.5 |
| Superior parietal lobe | R | | 3.51 | 29.42 | -46.38 | 57.25 |
| Operculum | R | 318 | 4.78 | 61.77 | -4.09 | 5 |
| Superior temporal gyrus | R | | 4.51 | 54.30 | -9.06 | -8.75 |
| Operculum | R | | 4.46 | 64.26 | -16.53 | 16 |
| Angular gyrus | R | | 4.24 | 61.77 | -38.92 | 10.5 |
| Posterior middle temporal gyrus | R | | 4.18 | 61.77 | -26.48 | -14.25 |
| Precentral gyrus | R | 243 | 4.68 | 31.91 | -6.58 | 49 |
| Precentral gyrus | R | | 4.16 | 39.38 | -9.06 | 51.75 |
| Precentral gyrus | R | | 3.93 | 39.38 | -19.02 | 43.5 |
| Precentral gyrus | R | | 3.67 | 54.30 | -4.09 | 49 |
| Precentral gyrus | R | | 3.46 | 31.91 | -9.06 | 68.25 |
| **Seed: Left SPL [-22.8, -71.3, 46.2]** | | | | | | |
| **FPM > WPM** | | | | | | |
| Posterior middle temporal gyrus | L | 473 | 5.20 | -60.14 | -56.34 | 7.75 |
| Posterior middle temporal gyrus | L | | 5.12 | -60.14 | -38.92 | 2.25 |
| Angular gyrus | L | | 4.60 | -40.24 | -58.82 | 21.5 |
| Lateral occipital cortex inferior | L | | 4.54 | -40.24 | -63.80 | 7.75 |
| Lateral occipital cortex inferior | L | | 4.18 | -55.17 | -66.29 | 7.75 |
| Superior parietal lobe | R | 349 | 4.45 | 39.38 | -48.87 | 49 |

| **Anatomical structure** | **Hemi** | ***k*** | ***t*** | ***x*** | ***y*** | ***z*** |
|---|---|---|---|---|---|---|
| Precentral gyrus | R | | 4.25 | 36.89 | -14.04 | 43.5 |
| Angular gyrus | R | | 4.07 | 41.86 | -58.82 | 46.25 |
| Supramarginal gyrus | R | | 3.99 | 54.30 | -38.92 | 54.5 |
| Superior parietal lobe | R | | 3.93 | 41.86 | -51.36 | 57.25 |
| **Seed: Left SPL [-22.8, -71.3, 46.2]** | | | | | | |
| **Tone > FPM** | | | | | | |
| Superior frontal gyrus | L | 436 | 6.78 | -20.34 | 35.72 | 29.75 |
| Frontal pole | L | | 4.63 | -22.82 | 55.62 | 27.00 |
| Frontal pole | L | | 4.54 | -20.34 | 43.18 | 18.75 |
| Middle frontal gyrus | L | | 4.42 | -37.75 | 38.21 | 32.50 |
| Frontal pole | L | | 4.15 | -15.36 | 53.14 | 32.50 |
| Frontal pole | R | 234 | 4.24 | 24.45 | 53.14 | 24.25 |
| Frontal pole | R | | 4.24 | 36.89 | 40.70 | 24.25 |
| Frontal pole | R | | 4.16 | 4.54 | 58.11 | 21.50 |
| Frontal pole | R | | 3.79 | 24.45 | 40.70 | 27.00 |
| Frontal pole | R | | 3.78 | 14.50 | 60.60 | 18.75 |
| **Seed: Left Occ. pole [-17.9, -91.2, 18.8]** | | | | | | |
| **FPM > WPM** | | | | | | |
| Cerebellum | L | 641 | 5.92 | -0.43 | -78.73 | -28 |
| Cerebellum | L | | 5.23 | -25.31 | -71.26 | -22.5 |
| Inferior temporal gyrus | L | | 4.84 | -52.68 | -53.85 | -14.25 |
| Fusiform gyrus | L | | 4.63 | -35.26 | -68.78 | -22.5 |
| Fusiform gyrus | L | | 4.53 | -10.38 | -78.73 | -19.75 |
| Lateral occipital cortex superior | L | 2134 | 5.92 | -17.85 | -83.70 | 27 |
| Intracalcarine cortex | R | | 5.44 | 4.54 | -86.19 | 7.75 |
| Superior parietal lobe | R | | 5.28 | 29.42 | -56.34 | 43.5 |
| Lateral occipital cortex superior | R | | 5.25 | 34.40 | -63.80 | 46.25 |
| Cuneal cortex | R | | 5.05 | 7.03 | -76.24 | 32.5 |
| Superior frontal gyrus | L | 172 | 5.21 | -22.82 | 25.77 | 54.5 |
| Middle frontal gyrus | L | | 4.79 | -25.31 | 28.26 | 38 |
| Frontal pole | L | | 3.57 | -17.85 | 48.16 | 43.5 |
| Frontal pole | L | | 3.45 | -27.80 | 38.21 | 43.5 |
| Middle frontal gyrus | L | | 3.39 | -40.24 | 18.30 | 51.75 |
| Precuneus | R | 766 | 5.11 | 9.52 | -48.87 | 49 |
| Superior parietal lobe | L | | 4.93 | -7.90 | -28.97 | 46.25 |
| Posterior cingulate cortex | R | | 4.83 | 12.01 | -36.43 | 35.25 |

| Anatomical structure | Hemi | *k* | *t* | *x* | *y* | *z* |
|---|---|---|---|---|---|---|
| Supplementary motor cortex | L | | 4.76 | -2.92 | -9.06 | 49 |
| Posterior cingulate cortex | L | | 4.55 | -10.38 | -38.92 | 40.75 |
| Precentral gyrus | R | 488 | 4.49 | 36.89 | -11.55 | 46.25 |
| Frontal pole | R | | 4.45 | 26.94 | 40.70 | 43.5 |
| Middle frontal gyrus | R | | 4.23 | 36.89 | 3.38 | 46.25 |
| Postcentral gyrus | R | | 4.11 | 36.89 | -21.50 | 43.5 |
| Middle frontal gyrus | R | | 4.06 | 26.94 | 28.26 | 49 |
| Superior temporal gyrus | R | 306 | 4.34 | 59.28 | 3.38 | -11.5 |
| Operculum | R | | 4.10 | 64.26 | -6.58 | 7.75 |
| Inferior frontal gyrus, pars opercularis | R | | 4.10 | 51.82 | 20.79 | 18.75 |
| Operculum | R | | 4.02 | 54.30 | -14.04 | 16 |
| Precentral gyrus | R | | 4.01 | 56.79 | 0.89 | 13.25 |
| Operculum | L | 188 | 4.23 | -55.17 | -6.58 | 7.75 |
| Posterior middle temporal gyrus | L | | 4.13 | -57.66 | -36.43 | -8.75 |
| Planum temporale | L | | 3.98 | -65.12 | -14.04 | 5 |
| Superior temporal gyrus | L | | 3.76 | -67.61 | -23.99 | 7.75 |
| Posterior middle temporal gyrus | L | | 3.69 | -67.61 | -33.94 | -3.25 |
| Cerebellum | R | 172 | 4.09 | 16.98 | -46.38 | -58.25 |
| Cerebellum | R | | 3.97 | 26.94 | -51.36 | -52.75 |
| Cerebellum | R | | 3.75 | 44.35 | -51.36 | -39 |
| Cerebellum | R | | 3.54 | 7.03 | -61.31 | -47.25 |
| Cerebellum | R | | 3.46 | 36.89 | -56.34 | -50 |
| **Seed: Left Intracalc. [-17.9, -66.3, 2.3]** | | | | | | |
| **FPM > WPM** | | | | | | |
| Superior temporal gyrus | L | 187 | 4.86 | -57.66 | -1.60 | -11.5 |
| Operculum | L | | 4.49 | -57.66 | -6.58 | 10.5 |
| Planum polare | L | | 4.40 | -55.17 | -1.60 | -3.25 |
| Insula | L | | 3.43 | -40.24 | -1.60 | 2.25 |
| Superior temporal gyrus | L | | 3.22 | -60.14 | -21.50 | -0.5 |

*Note.* Results are thresholded at $p < 0.01$ at peak level and FWE-corrected at $p < 0.05$ at cluster level.
